# PROX1 inhibits PDGF-B expression to prevent myxomatous degeneration of heart valves

**DOI:** 10.1101/2023.05.10.540284

**Authors:** Yen-Chun Ho, Xin Geng, Anna O’Donnell, Jaime Ibarrola, Amaya Fernandez-Celis, Rohan Varshney, Kumar Subramani, Zheila J Azartash-Namin, Jang Kim, Robert Silasi, Jill Wylie-Sears, Zahra Alvandi, Lijuan Chen, Boksik Cha, Hong Chen, Lijun Xia, Bin Zhou, Florea Lupu, Harold M. Burkhart, Elena Aikawa, Lorin E. Olson, Jasimuddin Ahamed, Natalia López-Andrés, Joyce Bischoff, Katherine E. Yutzey, R. Sathish Srinivasan

**Affiliations:** Cardiovascular Biology Research Program, Oklahoma Medical Research Foundation, 825 NE 13th street, Oklahoma City, OK 73104, USA; Division of Molecular Cardiovascular Biology, Cincinnati Children’s Hospital Medical Center, 3333 Burnet Ave, Cincinnati, OH 45229, USA; Molecular Cardiology Research Institute, Tufts Medical Center, Boston, MA; Cardiovascular Translational Research, Navarrabiomed (Miguel Servet Foundation), Instituto de Investigación Sanitaria de Navarra (IdiSNA), Hospital Universitario de Navarra (HUN), Universidad Pública de Navarra (UPNA), Pamplona, Spain; Department of Cell Biology, University of Oklahoma Health Sciences Center, Oklahoma City, OK 73117, USA; Vascular Biology Program, Boston Children’s Hospital, Boston, MA 02115, USA; Department of Genetics, Albert Einstein College of Medicine, Bronx, NY 10461, USA; Oklahoma Children’s Hospital, University of Oklahoma Health Heart Center, Oklahoma City, OK 73104, USA; Department of Medicine, Cardiovascular Division, Brigham and Women’s Hospital, Boston, MA 02115, USA

## Abstract

**Background:** Cardiac valve disease (CVD) is observed in 2.5% of the general population and 10% of the elderly people. Effective pharmacological treatments are currently not available, and patients with severe CVD require surgery. PROX1 and FOXC2 are transcription factors that are required for the development of lymphatic and venous valves. We found that PROX1 and FOXC2 are expressed in a subset of valvular endothelial cells (VECs) that are located on the downstream (fibrosa) side of cardiac valves. Whether PROX1 and FOXC2 regulate cardiac valve development and disease is not known.

**Methods:** We used histology, electron microscopy and echocardiography to investigate the structure and functioning of heart valves from *Prox1^ΔVEC^* mice in which *Prox1* was conditionally deleted from VECs. Isolated valve endothelial cells and valve interstitial cells were used to identify the molecular mechanisms *in vitro*, which were tested *in vivo* by RNAScope, additional mouse models and pharmacological approaches. The significance of our findings was tested by evaluation of human samples of mitral valve prolapse (MVP) and aortic valve insufficiency.

**Results:** Histological analysis revealed that the aortic and mitral valves of *Prox1^ΔVEC^* mice become progressively thick and myxomatous. Echocardiography revealed that the aortic valves of *Prox1^ΔVEC^* mice are stenotic. FOXC2 was downregulated and platelet-derived growth factor-B (PDGF-B) was upregulated in the VECs of *Prox1^ΔVEC^* mice. Conditional knockdown of FOXC2 and conditional overexpression of PDGF-B in VECs recapitulated the phenotype of *Prox1^ΔVEC^* mice. PDGF-B was also increased in mice lacking FOXC2 and in human MVP and insufficient aortic valve samples. Pharmacological inhibition of PDGF-B signaling with imatinib partially ameliorated the valve defects of *Prox1^ΔVEC^* mice.

**Conclusion:** PROX1 antagonizes PDGF-B signaling partially via FOXC2 to maintain the extracellular matrix composition and prevent myxomatous degeneration of cardiac valves.

**Novelty and Significance:** **What Is Known?**

- The transcription factors PROX1 and FOXC2 are critical regulators of lymphatic and venous valve development.
- PROX1 and FOXC2 are expressed in the downstream valvular endothelial cells of heart valves.

**What Is New?**

- Deletion of *Prox1* from the valvular endothelial cells of mice results in enlarged and myxomatous aortic and mitral valves. Aortic valves of the mutant (*Prox1^ΔVEC^*) mice were stenotic.
- FOXC2 is partially responsible for the phenotype of *Prox1^ΔVEC^* mice.
- PROX1 and FOXC2 inhibit the expression of the cytokine PDGF-B in heart valves.
- Hyperactivation of PDGF-B signaling results in aortic and mitral valve thickening.
- Inhibition of PDGF-B signaling ameliorates aortic valve stenosis in *Prox1^ΔVEC^* mice.
- *PDGFB* is overexpressed and *PROX1* is downregulated in human mitral valve prolapse (MVP) samples.

Our findings suggest that PROX1 is an inhibitor of myxomatous valve disease that afflicts ~10% of the elderly population. We have also identified PDGF-B as a potential target for treating myxomatous valve disease.

## INTRODUCTION

Blood flow into and away from the heart is regulated by the four heart valves. Healthy valves open and close approximately 40 million times a year, permitting blood flow in when they are open, and preventing regurgitation when they are closed. During each cardiac cycle, they experience shear stress while open due to blood flow, tension while closed due to withstanding the backflow, and flexure during opening and closing.

Heart valve disease is observed in ~2.5% of all individuals in the US, increasing to ~8% in individuals >65 years old and ~13% in those who are >75 years old ^1^. The rapid increase in the elderly population is expected to raise the frequency of heart valve disease. Valve repair surgery and prosthetic valves have significantly reduced the mortality associated with valve disorders; however, non-invasive approaches to stop or slow the progression of these diseases are urgently needed. The development of non-invasive treatments is hindered by our incomplete understanding of the pathological mechanisms underlying valve diseases.

Heart valve disorders are all associated with defects in the extracellular matrix (ECM). The rapid and reversible deformations of valve leaflets during the cardiac cycle depend on proper organization of the ECM. Heart valves contain three stratified layers of ECM: collagen-rich fibrosa on the outflow (downstream) side, proteoglycan-rich spongiosa in the middle and an elastin-rich layer on the inflow (upstream) side ^2^. Heart valves contain valvular interstitial cells (VICs) and valvular endothelial cells (VECs); the VICs are within the ECM and regulate its organization. A single layer of VECs surrounds VICs and the ECM on each side of the valve. VECs maintain the integrity of valves, sense shear stress and regulate ECM by secreting signaling molecules such as WNT9B, TGF-β and nitric oxide (NO) ^3–5^. Interestingly, VECs on the outflow and inflow sides of the valves display striking differences in gene expression ^6^, but the significance of VEC heterogeneity in VIC regulation and disease pathogenesis is unknown.

The homeobox transcription factor PROX1 is essential for the development of lymphatic, venous and lymphovenous valves ^7, 8^. We recently discovered that PROX1 is expressed in a subset of VECs that are located on the outflow side of heart valves ^8, 9^. The lymphedema-associated transcription factor FOXC2 is a target of PROX1 in vascular valves, and it is also expressed in the PROX1^+^ cells of heart valves ^10,11^. Wnt/β-catenin signaling regulates the expression of PROX1 and FOXC2 in both vascular and heart valves, suggesting that vascular and heart valves have shared regulatory mechanisms ^10^. Here, we use mouse models and human disease samples to test the hypothesis that PROX1 regulates the integrity of heart valves. We found that loss of *Prox1* in VECs during development results in the downregulation of FOXC2, heart valve thickening and myxomatous degeneration, resembling mitral valve prolapse (MVP) in humans. Furthermore, using loss-of-function and gain-of-function mouse models we have found that FOXC2 is partially responsible for the phenotype of mice lacking PROX1. Our work suggests that PROX1 and FOXC2 inhibit PDGF-B signaling and, ultimately, ECM organization in heart valves. Aberrant PDGF-B signaling was also observed in patients with MVP and aortic valve insufficiency. Our study suggests that PROX1 modulates myxomatous valve disease in humans, and that PDGF-B is a potential therapeutic target for treating valve disease.

## RESULTS

### PROX1 prevents the abnormal thickening of heart valves

We previously showed that PROX1 is expressed in a subset of CD31^+^ VECs that were located on the downstream side (fibrosa) of aortic valves at embryonic day (E) 16.5 ^8^. We analyzed embryos at earlier time points and found that PROX1 was not expressed in the atrio-ventricular cushion of E10.5 embryos (Figure S1A). However, PROX1 was expressed in the VECs of heart valves at least as early as E12.5 (Figure S1A). At E12.5 PROX1 expression did not appear to be obviously polarized. However, PROX1 expression became progressively restricted to the downstream VECs at subsequent time points and remained so in adult valves (Figure S1A). We generated a new *Prox1-2A-Cre* mouse model which express Cre recombinase from the regulatory elements of *Prox1* without compromising PROX1 expression (Figure S2A). We performed lineage tracing using this Cre line and determined that in postnatal day (P)7 *Prox1-2A-Cre*; *mT/mG* pups both upstream and downstream VECs were GFP^+^. This observation confirmed that PROX1 is initially expressed in both the upstream and downstream VECs. Additionally, we observed only a few GFP^+^ VICs in the *Prox1-2A-Cre*; *mT/mG* pups (Figure S1B). VICs originate from VECs through endothelial to mesenchymal transition (EndMT) before E10.5 ^12^. Hence, our lineage tracing result indicates that PROX1 expression in the heart valves starts at the conclusion of EndMT.

*Prox1^−/−^*embryos died at E14.5 due to the absence of lymphatic vasculature. At E14 the aortic valves of *Prox1^−/−^* embryos were indistinguishable from those in their wildtype littermates (Figure S3). Thus, PROX1 is not necessary for EndMT and the subsequent formation of heart valve leaflets.

To study the potential role of PROX1 in heart valve development, we aimed to delete *Prox1* specifically from VECs by using the *Prox1^f/f^* mice without affecting the lymphatic vasculature (Figure S2B) ^13^. To this end, we first used NFATC1^enCre^ that is expressed specifically in the downstream VECs of heart valves (Figure S2G) ^14^. Analysis of the aortic valves of postnatal day (P)7 NFATC1^enCre^; *Prox1^f/f^* pups revealed that the deletion of *Prox1* was incomplete as indicated by the residual PROX1 expression (Figure S4A, B). Nevertheless, the aortic valves of 6-month-old NFATC1^enCre^; *Prox1^f/f^* mice showed a trend towards thickening (Figure S4C-E).

We searched for alternative Cre lines to efficiently delete *Prox1* from VECs and investigate the valve phenotype thoroughly. In the *Notch1-Cre^Lo^*mice Cre recombinase replaces the Notch1 receptor intracellular domain at one *Notch1* locus ^15^. Cre is nuclear and active in tissues with high Notch signaling, such as in arteries and heart valves, but not in veins ^15^. We hypothesized that *Notch1-Cre^Lo^* will not be active in the lymphatic vasculature, as it originates predominantly from the embryonic veins. As anticipated, lineage tracing revealed that *Notch1-Cre^Lo^* was active in VECs and a substantial number of VICs (Figure S5A), but not in the lymphatic vessels of the mesentery or skin, and venous valves (Figure S5B-D).

Although heterozygous loss-of-function mutations in *NOTCH1* are associated with bicuspid aortic valves and calcific aortic valve disease in humans ^16^, several reports have shown that the heart valves of *Notch1^+/-^* mice are phenotypically normal ^5,17–19^. Consistent with these reports, 6-month-old *Notch1-Cre^Lo^* mice, which are heterozygous for *Notch1,* had phenotypically normal cardiac valves as evaluated by histological quantification of valve thickness (Figure S6A). *Notch1-Cre^Lo^* mice also showed normal aortic valve function at 12-months of age (Figure S6B).

*Notch1-Cre^Lo^; Prox1^f/f^* mice (referred to as *Prox1^ΔVEC^* for *Prox1* deletion in VECs) were born at the expected Mendelian ratio and did not display obvious lymphatic vascular defects such as edema, chylous ascites or chylothorax. GFP is expressed from the *Prox1* locus once the floxed allele is deleted by Cre recombinase ^13^. Compared to controls, *Prox1^ΔVEC^*mice expressed GFP in VECs and had significantly reduced numbers of PROX1-positive VECs in the aortic and mitral valves (Figure S7A, B), consistent with loss of *Prox1* in VECs. Notably, *Prox1^ΔVEC^* mice did not display bicuspid aortic valves (Figure S7A). Furthermore, analysis of the heart valves of *Prox1^ΔVEC^* mice by qRT-PCR and western blotting did not reveal any obvious changes in the expression of receptors, ligands or targets of Notch signaling (Figure S7C, D). Thus, Notch signaling is not affected in an obvious manner in the *Prox1^ΔVEC^* mice.

To examine heart valve thickness in adult mice, we stained sections with Movat pentachrome (Figure 1A). *Prox1^ΔVEC^* animals started to display thickened aortic and mitral valves when they were 3-month-old and their valves were significantly thicker than those from wild type, *Notch1-Cre^Lo^* or *Notch1-Cre^Lo^; Prox1^+/f^*littermates (henceforth combined as controls), at 6- and 12-months of age (Figure 1A and B). We reanalyzed the data from 6-month-old mice and determined that both male and female *Prox1^ΔVEC^* mice develop enlarged valves (Figure S8). Ultrasound image analysis revealed that the cusp separation and fractional valve opening of the aortic valves were reduced in 12-month-old *Prox1^ΔVEC^* mice when compared with the controls. Furthermore, color Doppler imaging revealed an increase in outflow velocity (Figure 1C and Video S1 and 2). These echocardiography studies indicated that the *Prox1^ΔVEC^* mice had aortic valve stenosis phenotype. Taken together, loss of PROX1 in VECs resulted in progressive thickening of aortic and mitral valves and aortic valve stenosis in a Notch signaling- and sex-independent manner.

**Figure 1:**
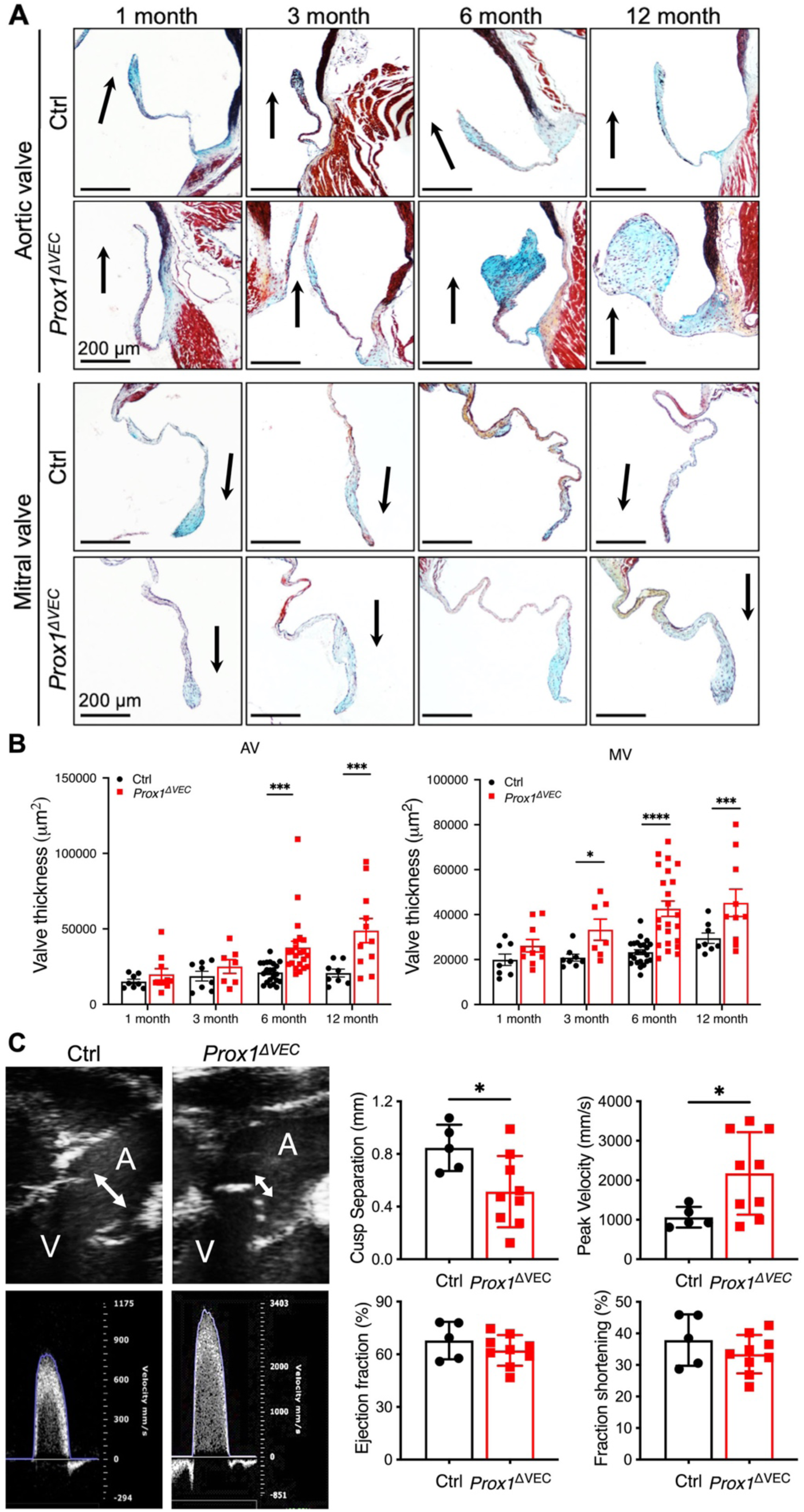
Deletion of *Prox1* from VECs results in progressive myxomatous valve degeneration and aortic valve stenosis. (A) Representative Movat Pentachrome-stained histologic images of the aortic and mitral valves of control and *Prox1^ΔVEC^* mice at various ages. Glycosaminoglycans, collagen and elastin are stained in blue, red and black, respectively. (B) Quantification of aortic valve and mitral valve thickness. For each of the mice four to seven sections at 100-120 μm intervals were analyzed. The area of the valve leaflets (2 leaflets from base to tip) was measured for every section and their average was used as a measure of valve thickness. Data is represented as Mean ± SEM. Statistical significance was assessed using a two-tailed, unpaired *t* test. Each dot represents an individual mouse, n=8-23 mice/genotype/time point. **** = *p* < 0.0001; *** = *p* < 0.001; ** = *p* < 0.01; * = *p* < 0.05. (C) Echocardiography of 12-month-old control and *Prox1^ΔVEC^* mice. Left: representative echocardiography images show decreased cusp separation (double-headed arrows in the top panels) and increased aortic peak velocity (bottom panels) in *Prox1^ΔVEC^* mice. Right: bar charts of cusp separation and aortic valve peak velocity. No obvious defects were observed in left ventricle functions in the *Prox1^ΔVEC^* mice as indicated by normal ejection fraction and fractional shortening. Data is represented as Mean ± SD. Statistical significance was assessed using a two-tailed, unpaired *t* test. n=5 control and n=9 *Prox1^ΔVEC^* mice. * = *p* < 0.05. Abbreviations: AV, aortic valve; MV, mitral valve; A, aorta; V, left ventricle.

### PROX1 inhibits late-onset myxomatous degeneration of heart valves

We performed scanning electron microscopy (SEM) to visualize the outflow side of the aortic and mitral valves. SEM confirmed that the *Prox1^ΔVEC^* mice do not develop bicuspid aortic valves (Figure 2A). In both control and mutant mice, VECs along the edges of the valves were elongated and those at the sites of coaptation had a cobblestone morphology (Figure 2A and Figure S9A). *Prox1^ΔVEC^* mice frequently showed disrupted cell junctions, detachment of the VEC layer, and platelet aggregate-like structures at the sites of coaptation in aortic valves (Figure 2A) but not in the mitral valves (Figure S9A). In Movat pentachrome-stained thin sections we observed platelet aggregate-like structures on the upstream (inflow) side of mutant aortic valves and in some mutant mitral valves (Figure 2B and Figure S9B). Additionally, the expression of von Willebrand Factor (vWF), a potent regulator of coagulation, was significantly increased in the aortic and mitral valves of *Prox1^ΔVEC^* mice (Figure 2C and Figure S9C). Both PROX1 and FOXC2 are known to inhibit thrombosis in venous valves ^20^. We analyzed the valves of *Prox1^ΔVEC^*mice and determined that the expression of FOXC2 was significantly downregulated (Figure 2D and Figure S9D). Thus, PROX1 regulates the expression of FOXC2 in VECs. Platelet aggregate-like structures are observed on both the downstream (PROX1^+^) and upstream (PROX1^-^) sides of cardiac valves, and these defects are more frequently observed in aortic valves. We infer that high shear stress on the tips and upstream side of abnormally thick *Prox1^ΔVEC^* aortic valves compromises VEC integrity.

**Figure 2:**
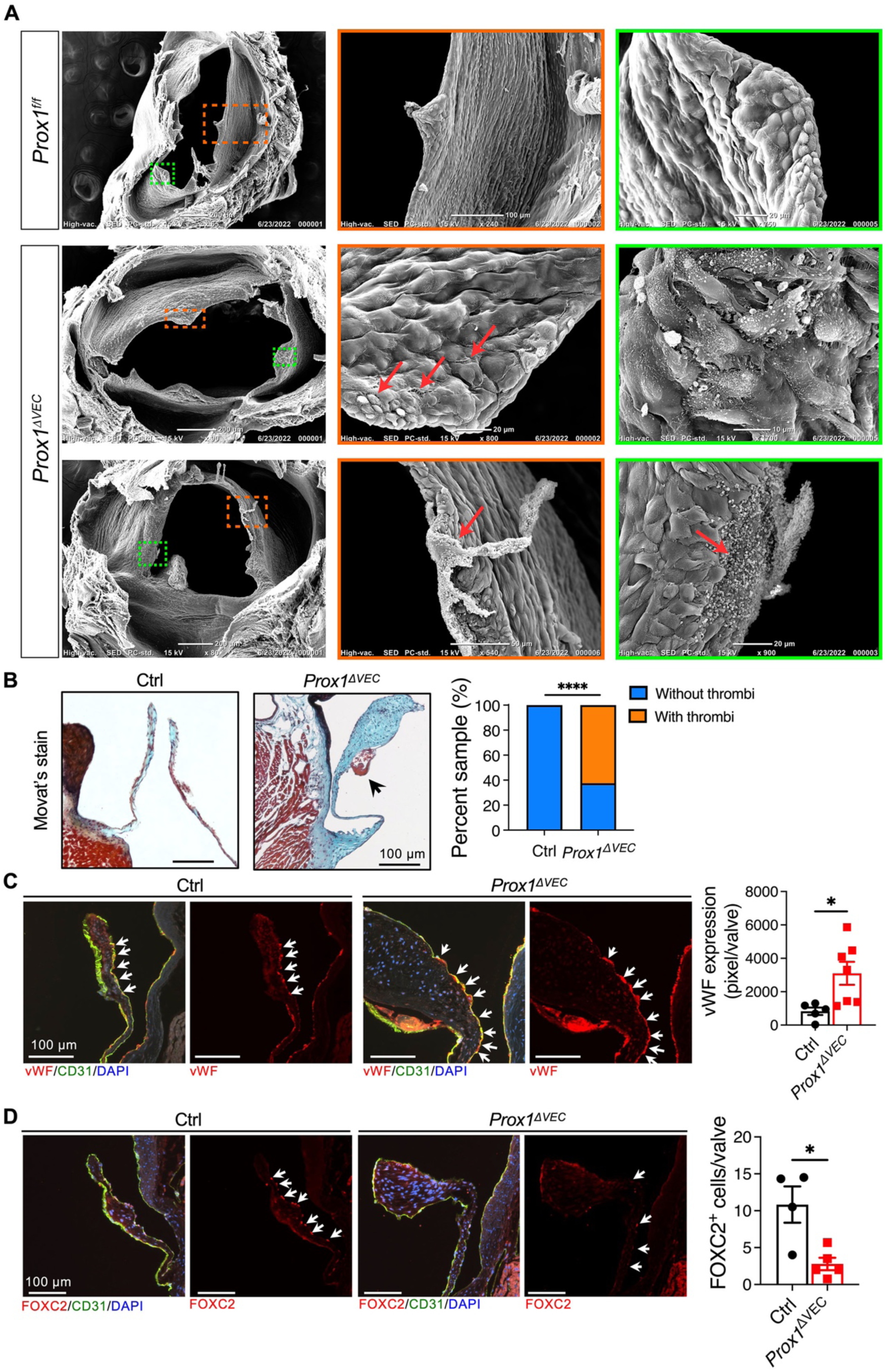
Loss of PROX1 from aortic VECs results in valve thickening, damaged endothelium, thrombus formation and downregulation of FOXC2 in VECs. (A) Representative scanning electron microscope (SEM) images of the tri-cuspid aortic valve leaflets of control and *Prox1^ΔVEC^* mice. SEM images show that *Prox1^ΔVEC^* mice have thicker aortic valves, disrupted endothelial layer (arrows in the middle row) with infiltration of platelet-like cells (arrows in the bottom row). N=5 for the control and N=6 for *Prox1^ΔVEC^*mice. (B) Representative Movat Pentachrome-stained histologic images demonstrated thrombus-like structure in the aortic valve of a *Prox1^ΔVEC^* mouse (arrow). The graph shows the quantification of the percentage of mice with thrombi in the aortic valves. Valves were analyzed by either SEM or Movat Pentachrome-staining of sections. N=10 control mice (5 by SEM, 5 by staining) and n=16 *Prox1^ΔVEC^* mice (6 by SEM and 10 by staining). (C) vWF expression was increased in the downstream side VECs of *Prox1^ΔVEC^* mice (arrows). N=5 for control mice; n=7 for *Prox1^ΔVEC^* mice. (D) FOXC2^+^ VECs (arrows) were reduced in *Prox1^ΔVEC^* mice. N=4 for control mice; n=5 for *Prox1^ΔVEC^*mice. Data is represented as Mean ± SEM. Statistical significance was assessed using the Fisher exact test in (B) and a two-tailed, unpaired *t* test in (C) and (D). * = *p* < 0.05; **** = *p* < 0.0001. Each dot represents the average of 3-5 sections from a mouse.

Abnormal ECM composition is a defining characteristic valve diseases ^2^. Hence, we investigated if the expression of collagen, elastin, and proteoglycans were altered in the thickened valves *Prox1^ΔVEC^* mice. Movat pentachrome-staining revealed that proteoglycan expression (blue staining) is increased in the valves of *Prox1^ΔVEC^* mice compared to controls (Figure 1A). Consistent with this observation we observed elevated expression of the proteoglycans aggrecan and versican in the aortic (Figure 3A, B) and mitral (Figure S10A, B) valves of *Prox1^ΔVEC^* mice.

**Figure 3:**
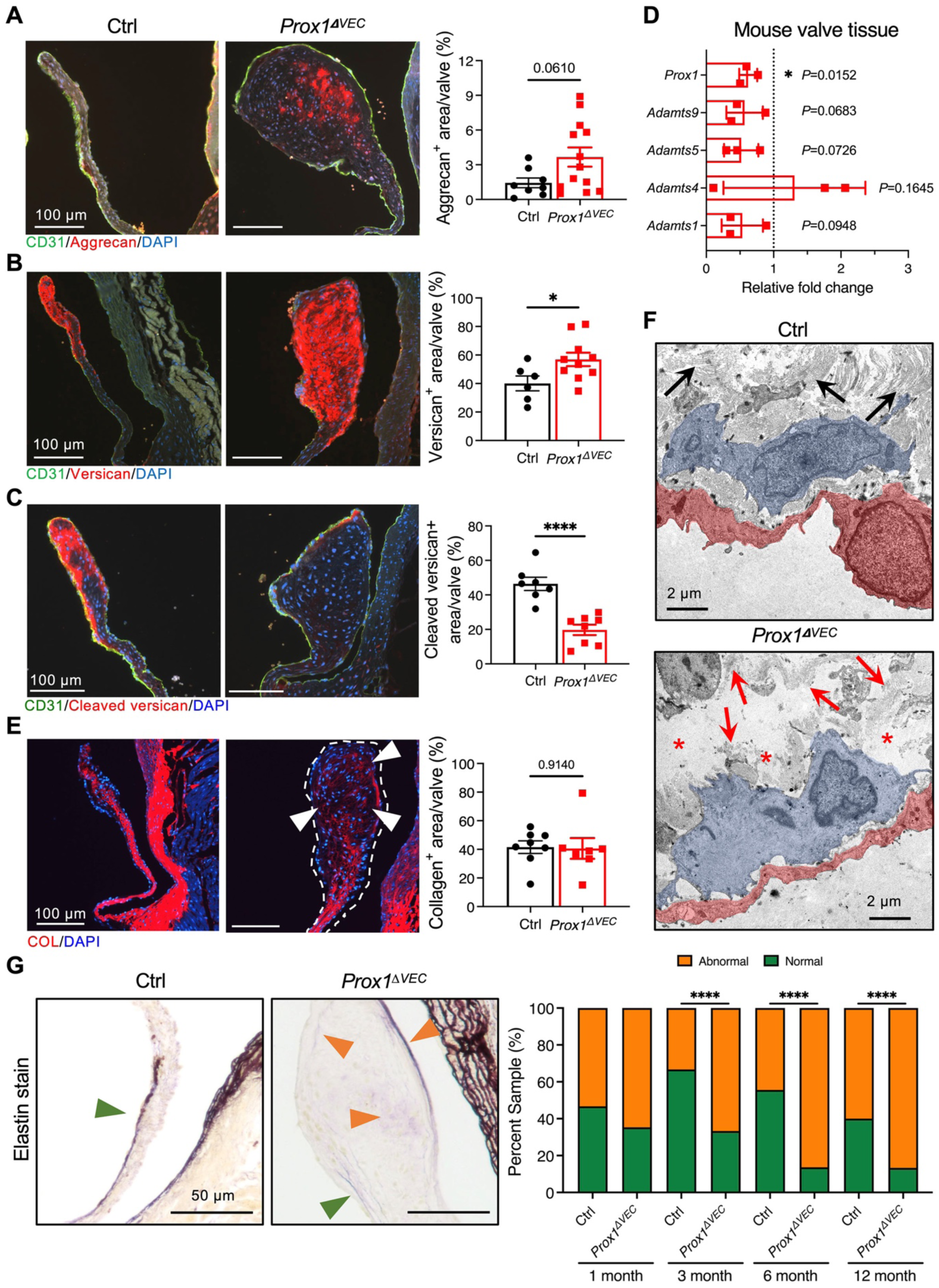
Increased proteoglycan expression and disrupted collagen and elastin fibers are observed in the aortic valves of *Prox1^ΔVEC^*mice. (A-C) Representative immunofluorescence images and semi-quantitative measurements of aggrecan (A), versican (B) and cleaved versican (anti-DPEAAE peptide in panel C) expression in the aortic valve sections from 6-month-old control and *Prox1^ΔVEC^*mice. Aggrecan and Versican were significantly increased and cleaved-versican was significantly reduced in *Prox1^ΔVEC^* mice. Each dot in the graph represents the average of 3-5 sections from a mouse. (D) RNA was extracted from aortic and mitral valve tissues pooled together from five 3-month-old control or five *Prox1^ΔVEC^* mice. N=3 control and n=3 *Prox1^ΔVEC^* pooled samples. qRT-PCR showed that 3 out of the 4-candidate proteoglycan degrading enzymes had a trend towards reduced expression in *Prox1^ΔVEC^* pooled samples. (E) Collagen binding peptide staining followed by semi-quantitative measurement suggested comparable expression levels in the aortic valves of 6-month-old control and *Prox1^ΔVEC^* mice. However, the collagen fibers in the aortic valves of *Prox1^ΔVEC^* mice appeared to be disrupted (arrowhead). N=8 control and n=7 *Prox1^ΔVEC^* mice (F) Representative transmission electron microscope (TEM) images show VECs (pseudocolored in brown) and VIC-like cells (pseudocolored in blue) in the aortic valves of 12-month-old mice. Well-organized collagen fibers (black arrows) were seen in control mice. In contrast, collagen fibers appeared to be disrupted in *Prox1^ΔVEC^* mice (red arrows) with gaps in between the bundles (red asterisk). (G) Resorcin-Fuchsin staining showed normal elastin expression in the upstream side of aortic valves (green arrowhead) in 6-month-old control mice. In contrast, normal elastin distribution was reduced and abnormal distribution within and in the downstream side of aortic valves (orange arrowhead) was observed in 6-month-old *Prox1^ΔVEC^* mice. The graph shows normal and abnormal elastin expression in control and *Prox1^ΔVEC^* mice at various ages. Data is represented as the percentage of total mice. 1-month-old mice: n=8 controls and n=9 *Prox1^ΔVEC^*; 3-month-old mice: n=7 controls and n=8 *Prox1^ΔVEC^*; 6-month-old mice: n=11 controls and *Prox1^ΔVEC^*; 12-month-old mice: n= 5 controls and n=8 *Prox1^ΔVEC^*. Data is represented as Mean ± SEM in (A, B, C, and E). Data is represented as Mean ± SD in (D). Statistical significance was assessed using a two-tailed, unpaired *t* test in (A, B, C, and E), using a one sample *t* and Wilcoxon test in (D), and the Fisher exact test in (G). * = *p* < 0.05, **** = *p* < 0.0001.

The ADAMTS family of proteases play an important role in degrading versican and aggrecan ^21^. Among the 19 members of the family, ADAMTS-1, -4, -5, -9, -15 and -20 can cleave versican and aggrecan ^21^. *Adamts-1, -4, -5* and *-9* are downregulated in human myxomatous mitral valves ^22^, and *Adamts9^+/-^* and *Adamts5^−/−^* mice develop myxomatous heart valves ^23,24^. Mutations in *ADAMTS19* are associated with non-syndromic progressive heart valve disease ^25^. We hypothesized that the elevated levels of aggrecan and versican in the valves of *Prox1^ΔVEC^* mice arises from reduced ADAMTS activity. To test this hypothesis, we used IHC to detect a peptide fragment (DPEAAE) that is released by ADAMTS-mediated cleavage of versican. Consistent with a previous report, we observed high levels of cleaved versican (DPEAAE) in control valves (Figure 3C and Figure S10C)^24^. In contrast, DPEAAE levels were significantly reduced in the aortic valves of 6-month-old *Prox1^ΔVEC^*mice (Figure 3C) and showed a trend towards reduction in the mitral valves of mutants (Figure S10C). We extracted RNA from aortic and mitral valves of 3-month-old control and *Prox1^ΔVEC^* mice and found that *Adamts1, 5,* and *9* transcript levels were reduced, albeit in a non-significant manner, in *Prox1^ΔVEC^* valves (Figure 3D).

IHC with collagen hybridizing peptide did not reveal significant changes in collagen expression in *Prox1^ΔVEC^* mice compared to controls (Figure 3E and Figure S10D). However, collagen fibers appeared to be disrupted. Indeed, collagen fibers appeared to be disrupted in portions of the aortic valves of *Prox1^ΔVEC^*mice in transmission electron microscopy (TEM) images as well (Figure 3F, red arrows). Correspondingly, collagen fibers appeared to be absent from portions of the valve (Figure 3F, asterisk). Elastin fibers, which are normally localized to the upstream side, were quantified as described by Gomez-Stallons et al ^26^. The elastin fibers were disrupted and mis-localized to the downstream and interstitial layers in *Prox1^ΔVEC^*valves (Figure 3G and Figure S10E).

We did not observe Alizarin red^+^ calcific nodules in the aortic valves of chow or high-fat diet fed *Prox1^ΔVEC^* mice (Figure S11). Thus, *Prox1^ΔVEC^* mice do not develop calcific aortic valve disease. Instead, deletion of *Prox1* from VECs led to the progressive thickening of heart valves, disruption of VEC layer with platelet infiltration, aberrant accumulation of unprocessed proteoglycans and disruption of collagen and elastin fibers. Based on the combined data, we conclude that *Prox1^ΔVEC^*valves undergo late-onset myxomatous degeneration ^27,28^.

### PDGF-B and FOXC2 are physiologically relevant target of PROX1 in VECs

VICs control ECM deposition and organization. Abnormal EndMT could result in an increase in VIC numbers and consequently ECM disorganization. We determined that the VIC numbers were not increased, and that their density was reduced in the aortic and mitral valves of *Prox1^ΔVEC^* mice (Figure S12). We also did not observe any obvious increase in the number of smooth muscle actin (SMA)^+^ myofibroblasts (a marker of EndMT) in 6-month-old *Prox1^ΔVEC^* mice although a mild to moderate increase was observed in 12-month-old mutants (Figure S13). These observations suggested that the ECM composition of *Prox1^ΔVEC^* mice is not defective due to an increase in the number of VICs or EndMT but is likely caused by an imbalance in ECM synthesis and degradation.

VECs are known to regulate VICs in a paracrine manner. To determine if signaling defects contribute to valve thickening in *Prox1^ΔVEC^*mice, we examined the abundance of transcripts encoding 12 common cytokines and growth factors. We found that *Platelet-derived growth factor B* (*Pdgfb*) and *transforming growth factor B1* (*Tgfb1*) were significantly elevated in *Prox1^ΔVEC^* valves compared to controls (Figure 4A), suggesting that PROX1 negatively regulates the expression of these genes in VECs. To test this hypothesis, we used a PROX1-specific siRNA in sheep mitral VECs, and observed upregulation of *PDGFB,* but not *TGFB1* (Figure S14). Moreover, we used the modified *in situ* hybridization approach RNAscope to validate the qRT-PCR data for *Pdgfb*. We observed a few *Pdgfb^+^*puncta in control valves. In contrast, *Pdgfb* was strikingly upregulated in both VECs and VICs of 3-month-old *Prox1^ΔVEC^* mice (Figure 4B and Figure S15A). These observations are consistent with the upregulation of *Pdgfb* in *Prox1^ΔVEC^* VECs.

**Figure 4:**
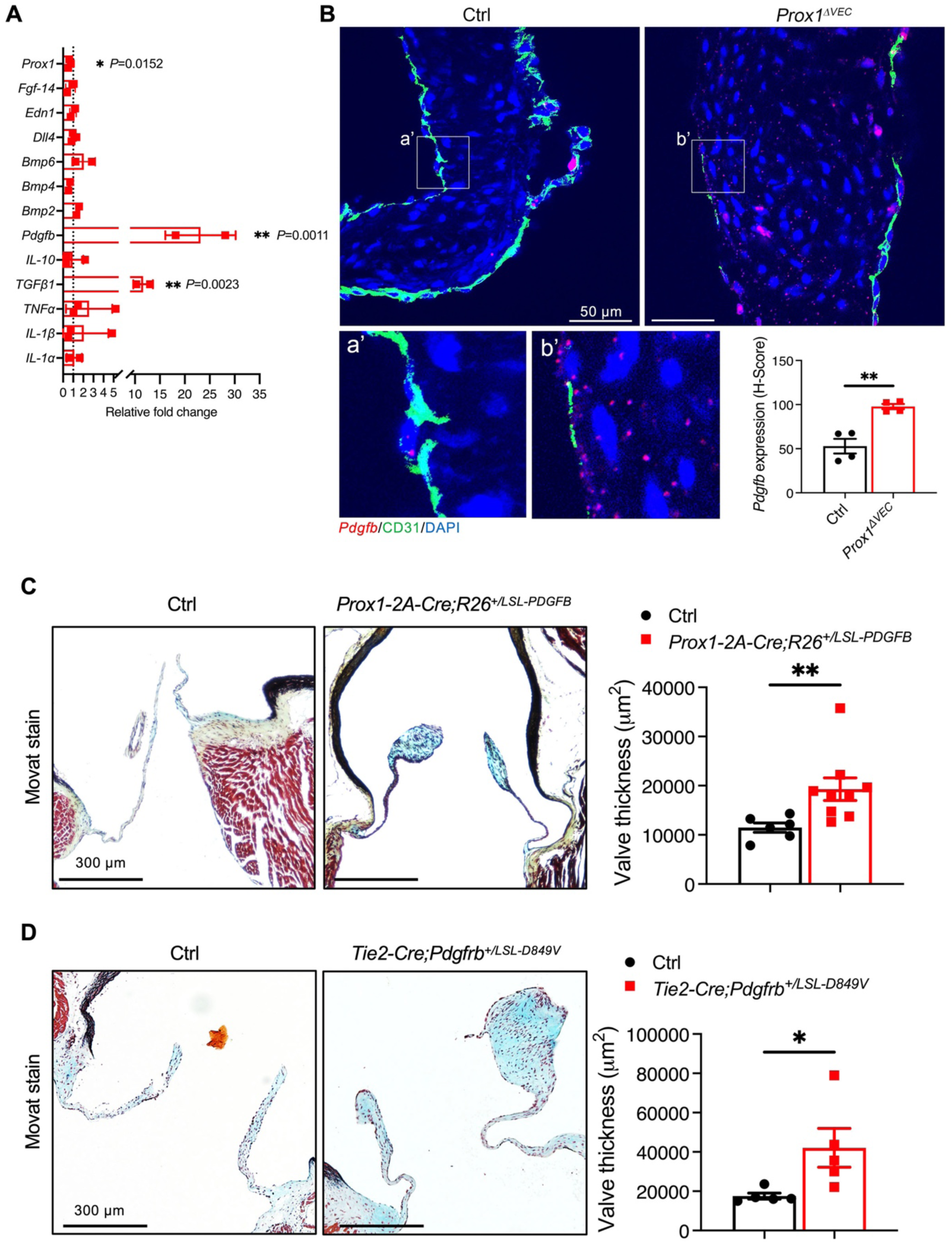
Hyperactivation of PDGF-B/PDGFRβ signaling causes myxomatous degeneration of aortic valve. (A) qRT-PCR for the expression of selected cytokines and growth factors in the valve tissue from control and *Prox1^ΔVEC^* mice. The dotted line represents normalized expression in control valves. Each dot represents aortic and mitral valve tissues collected from five 3-month-old mice. Data is represented as Mean ± SD. (B) Representative RNAscope images for *Pdgfb* expression (red dots) in the aortic valves of 3-month-old control and *Prox1^ΔVEC^* mice. The lower panels are enlarged images of the boxed areas in the upper panels. Quantification of the RNAscope results showed that *Prox1^ΔVEC^* mice had higher *Pdgfb* expression. Each dot represents the average from 3 sections of a mouse valve. N=4 mice per genotype. (C) Representative Movat Pentachrome-stained images of aortic valves from 6-month-old control and *Prox1-2A-Cre;R26^+/LSL-PDGFB^* mice. Thicker aortic valves were observed in *Prox1-2A-Cre;R26^+/LSL-PDGFB^* mice when compared with control valves. N=6 control and n=9 *Prox1-2A-Cre;R26^+/LSL-PDGFB^* mice. (D) Representative Movat Pentachrome stain images of the aortic valves of 6-month-old control and Tie2-Cre*;Pdgfrb^+/D849V^* mice. Tie2-Cre*;Pdgfrb^+/D849V^*mice had enlarged aortic valves when compared to control valves. N=5 for control and Tie2-Cre*;Pdgfrb^+/D849V^* mice. The area of valve leaflets was measured from 4-6 stained sections per valve. The average of these areas is represented as the thickness of valve in that mouse. Statistical significance was assessed using a two-tailed, unpaired *t* test in (B, C, and D) and using a one sample *t* and Wilcoxon test in (A). * = *p* < 0.05. ** = *p*< 0.01.

TGF-β1 is a known regulator of ECM composition, and mutations in genes that regulate TGF-β signaling are associated with syndromic mitral valve defects such as Marfan syndrome ^29^. PDGF-B is also a potent regulator of ECM composition and *Pdgfb^−/−^* mice that die perinatally possess hypoplastic heart valves ^30,31^. However, little is known about the role of PDGF-B in heart valve disease. We bred the *Prox1-2A-Cre* and *R26^+/LSL-PDGFB^* mice to overexpress PDGF-B in the VECs (Figure S2A, C, G) ^32^. The aortic and mitral valves of *Prox1-2A-Cre* and *R26^+/LSL-PDGFB^* mice were significantly thicker than that of their littermates (Figure 4C and Figure S15B). This data suggested that the increased expression of PDGF-B in VECs is responsible for the thickening of *Prox1^ΔVEC^*valves.

PDGF receptor β (PDGFRβ) is the cognate receptor of PDGF-B, and it is expressed in connective tissue cells such as VICs. *De novo* W566R or P584R mutations in PDGFRβ result in constitutive activation of the receptor and the rare disease Kosaki overgrowth syndrome ^33,34^. Mitral Valve Prolapse (MVP) was identified in a patient with the W566R mutation and in another patient with the P584R mutation ^33,35^. To investigate the physiological significance of PDGF-B signaling in valve disease, we used a previously reported model to conditionally express constitutively active PDGFRβ^D849V^ in VICs ^36^. The D849V mutation is in the kinase domain of the receptor. We bred the *Pdgfrb^+/LSL-D^*^849*V*^ with *Tie2-Cre* to induce PDGFRβ^D849V^ expression specifically in the PDGFRβ^+^ cells of heart valves (Figure S2D, G) ^12^. Analysis of 6-month-old mice revealed significantly thicker valves and more proteoglycan deposition (blue staining) in *Tie2-Cre; Pdgfrb^+/LSL-D849V^*mice compared to control littermates (Figure 4D and Figure S15C). Thus, overexpression of PDGF-B in VECs or hyperactivation of PDGFRβ signaling in VICs can recapitulate the phenotype of *Prox1^ΔVEC^* mice.

Together, our results suggest that PDGF-B is a physiologically relevant target of PROX1 and a novel regulator of myxomatous valve disease. PROX1 could directly regulate PDGF-B or indirectly through an intermediate transcription factor. As shown in Figure 2D and Figure S9D FOXC2 expression is downregulated in the VECs of *Prox1^ΔVEC^* mice. FOXC2 inhibits PDGF-B expression in the lymphatic endothelial cells ^37,38^. Hence, we wanted test if FOXC2 is a physiologically relevant target of PROX1 in VECs. *Foxc2^−/−^* embryos die between E13.5 and birth. We have developed a novel shRNA-based strategy to overcome the embryonic lethality of *Foxc2^−/−^* mice and to knockdown *Foxc2* in VECs (Figure S2E)^39^. We combined *Notch1-Cre^Lo^* and CAG-LoxP-Stop-LoxP-rtTA3-mKate2 mice to express reverse tetracycline transactivator 3 (rtTA3) and the fluorescent marker mKate2 in VECs. In the presence of doxycycline (Dox), rtTA3 binds and activates the tetracycline response element (TetO), inducing expression of GFP and a highly efficient, miRNA-based shRNA targeting *Foxc2* (Figure S2E).

We administered Dox to *Notch1-Cre^Lo^; CAG-LoxP-Stop-LoxP-rtTA3-mKate2; TetO-GFP-shFoxc2* mice (henceforth called as *shFoxc2*) from E10.5 and analyzed the heart valves at 12 months of age. Our analysis revealed that the aortic valves of *shFoxc2* mice were significantly thicker than that of control littermates (Figure 5A). Additionally, elastin expression was defective, the proteoglycan versican was enriched, and *Pdgfb* expression was increased in the aortic valves of *shFoxc2* mice (Figure 5B-D).

**Figure 5:**
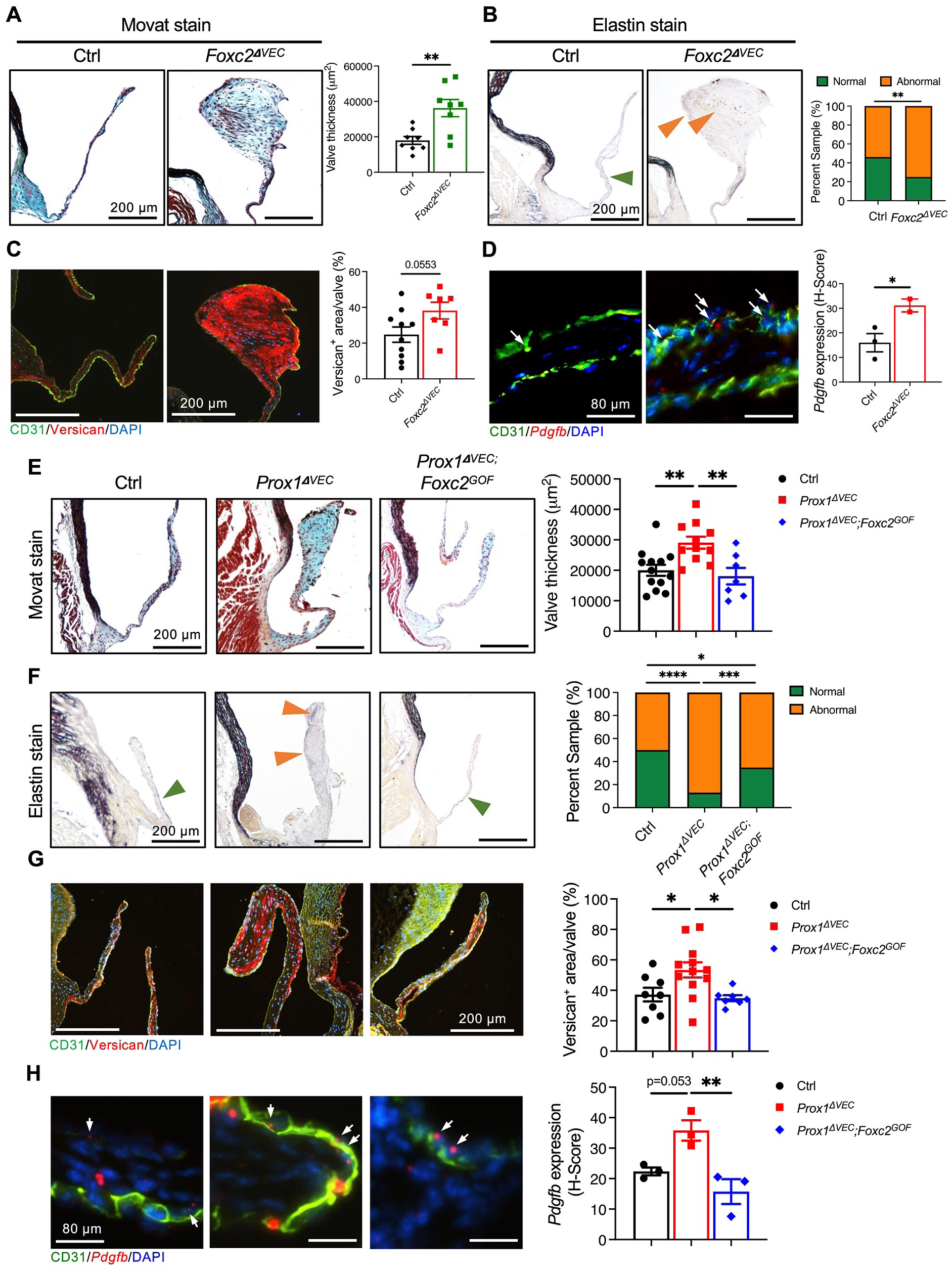
Knock down of FOXC2 results in enlarged aortic valves, and over expression of FOXC2 rescues the aortic valve defects of *Prox1^ΔVEC^* mice. (A) Representative images of Movat Pentachrome staining that was performed using aortic valves of 12-month-old control and *Foxc2^ΔVEC^* mice with graph showing the semi-quantitative measurement of valve thickness. N=8 controls and n=8 *Foxc2^ΔVEC^*mice. (B) Representative images of Resorcin-Fuchsin staining that was performed using the aortic valves of 12-month-old control and *Foxc2^ΔVEC^* mice. Green arrowhead shows the normal expression of elastin on the upstream side of valves. The orange arrowheads show abnormal elastin expression within and in the downstream side of valves. The graph shows the distribution of normal and abnormal elastin expression in control and mutant mice. N=11 controls and n=7 *Foxc2^ΔVEC^*mice. (C) Representative immunofluorescence images for versican expression in the aortic valves of 12-month-old control and *Foxc2^ΔVEC^*mice, followed by semi-quantitative measurement. N=10 control and n=8 *Foxc2^ΔVEC^* mice. (D) Representative RNAscope images for *Pdgfb* expression (red dots) in the aortic valves of 6-month-old control and *Foxc2^ΔVEC^* mice (arrows indicate *Pdgfb*^+^ VECs). Quantification of the RNAscope results showed that *Foxc2^ΔVEC^* mice had higher *Pdgfb* expression. Each dot represents the average of 3 sections from a mouse valve. N=3 control and n=2 *Foxc2^ΔVEC^* mice. (E-H) Over expression of FOXC2 in *Prox1^ΔVEC^* mice (*Prox1^ΔVEC^;Foxc2^GOF^*) ameliorates valve thickness (E), abnormal elastin distribution (F), increased proteoglycan expression (G) and *Pdgfb* expression (H). N=13 control, n=11 *Prox1^ΔVEC^* and n=7 *Prox1^ΔVEC^;Foxc2^ΔGOF^*mice in (E); N=13 control, n=11 *Prox1^ΔVEC^* and n=9 *Prox1^ΔVEC^;Foxc2^ΔGOF^*mice in (F); N=8 control, n=12 *Prox1^ΔVEC^* and n=7 *Prox1^ΔVEC^;Foxc2^ΔGOF^* mice in (G). N=3 for all three genotypes in (H). Mice were 6-month-old in (E-G) and 3-month-old in (H). Data is represented as Mean ± SEM. Statistical significance was assessed using a two-tailed, unpaired *t* test in (A, C, and D), one-way ANOVA and Tukey’s multiple comparisons test in (E, G, and H), and Fisher exact test in (B and F). * = *p* < 0.05; ** = *p* < 0.01; *** = *p* < 0.001; **** = *p* < 0.0001. Each dot represents the average of 3-5 sections from a mouse.

We previously reported a transgenic mouse model to overexpress FOXC2 in a Cre-dependent manner (Figure S2E) ^10^. In these FOXC2^GOF^ transgenic mice FOXC2 cDNA was inserted downstream of the ubiquitously expressed CMV β-actin promoter and a LoxP-GFP-Stop-LoxP cassette ^10^. FOXC2 is expressed in cells that express Cre and their progeny.

By breeding FOXC2^GOF^ with *Notch1-Cre^Lo^* we generated FOXC2^GOF^; *Prox1^ΔVEC^*mice to test whether restoration of FOXC2 expression could rescue the phenotype of *Prox1^ΔVEC^* mice. Indeed, the thickness of the aortic valve is reduced in FOXC2^GOF^; *Prox1^ΔVEC^* compared to *Prox1^ΔVEC^* mice (Figure 5E). Furthermore, normal expression pattern of elastin was restored and the expression of versican and *Pdgfb* was normalized in the FOXC2^GOF^; *Prox1^ΔVEC^* mice (Figure 5F-H).

The mitral valves of *shFoxc2* mice were not obviously defective (Figure S16A, B) and restoration of FOXC2 expression was not sufficient to ameliorate the mitral valve defects of *Prox1^ΔVEC^* mice (Figure S16C, D). Together these data suggested that FOXC2 is a physiologically relevant target of PROX1 in aortic valves. Yet to be identified targets of PROX1 are likely compensating for the loss of FOXC2 in mitral valves.

### PDGF-B upregulates proteoglycan expression in a SOX9-dependent manner

Valves share numerous similarities with connective tissue cell types such as chondrocytes ^40^, and constitutive activation of PDGFRβ upregulates the transcription factor SOX9 in chondrocytes ^30^. SOX9 is also upregulated in myxomatous valves from human and mice ^41,42^. Hence, we asked if SOX9 is regulated by PROX1 and PDGF-B in heart valves. We found that SOX9 was expressed in a few aortic VECs (Figure 6A, arrows) and weakly in the VICs (Figure 6A, arrowheads) of adult control mice as previously reported ^43^. In contrast, SOX9 was strongly upregulated in the VICs of *Prox1^ΔVEC^* mice (Figure 6A). SOX9 expression was also significantly upregulated in the aortic VICs of 6-month-old *Tie2-Cre; Pdgfrb^+/LSL-D849V^* mice (Figure 6B). Consistent with these observations, SOX9 was also overexpressed in the mitral VICs of *Prox1^ΔVEC^* and *Tie2-Cre; Pdgfrb^+/LSL-D849V^* mice (Figure S17A, B). Together, these findings are consistent with the hypothesis that PROX1 negatively regulates PDGFR-B signaling and, in turn, SOX9 expression.

**Figure 6:**
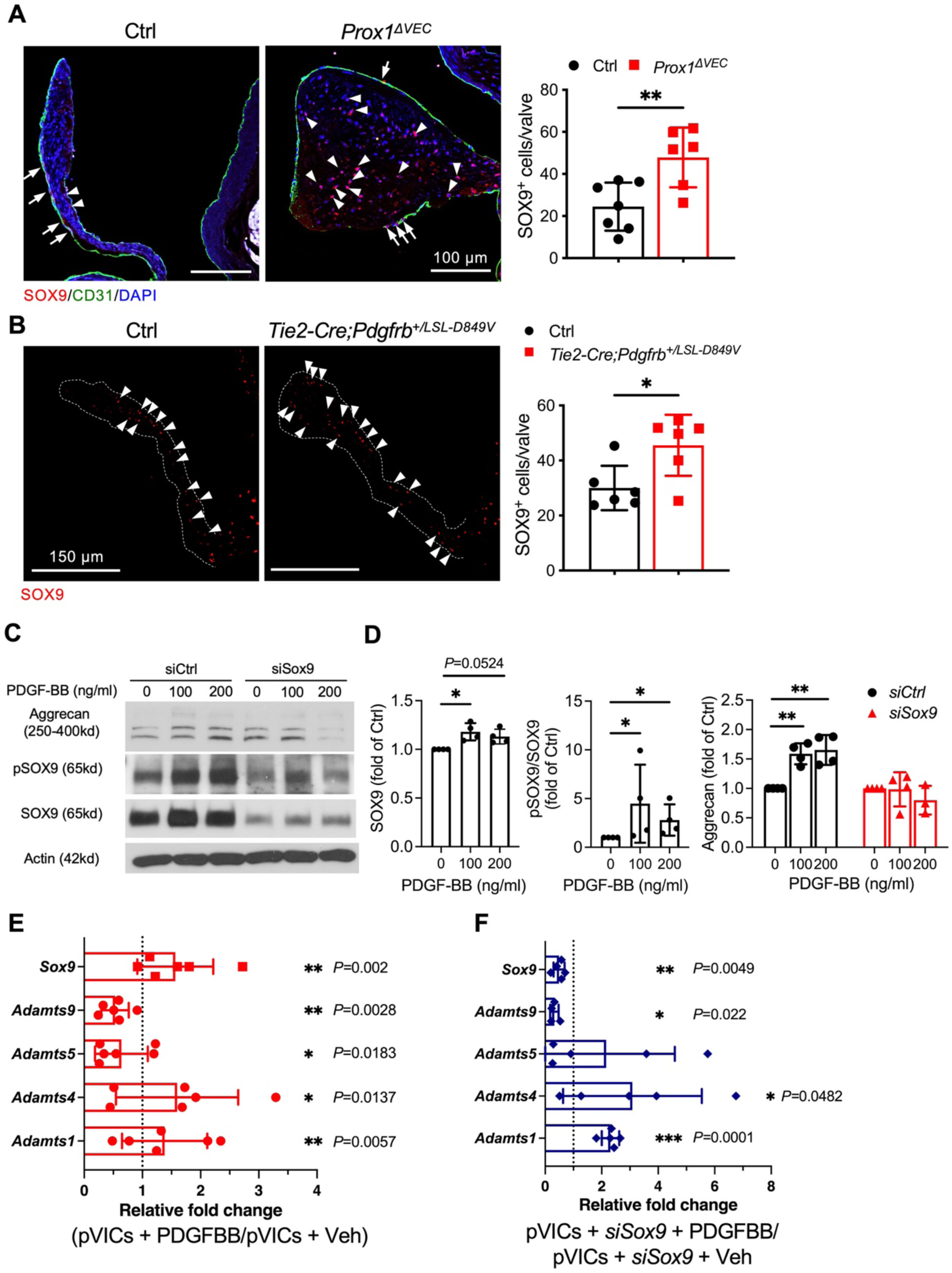
PDGF-B/PDGFRB signaling promotes proteoglycan expression in VICs in a SOX9-dependent manner. (A) Representative IHC images and quantification showing increased number of SOX9^+^ cells in the of aortic valves 6-month-old *Prox1^ΔVEC^* mice when compared to control littermates. Arrows indicate SOX9^+^ VECs and arrowheads indicate SOX9^+^ VICs. N=7 control and n=6 *Prox1^ΔVEC^* mice. (B) SOX9 expression and quantification in the valves of 6-month-old control and *Tie2-Cre;Pdgfrb^+/D849V^*mice. Arrowheads indicate SOX9^+^ VICs. N=7 for control and *Tie2-Cre;Pdgfrb^+/D849V^*mice. (C) Representative western blotting images using lysates prepared from porcine valve interstitial cells (pVICs) transfected with siControl or siSOX9 and treated with the indicated amount of PDGF-B for 24 hrs. (D) Quantification of the western blotting data shows that PDGF-B increases the expressions of aggrecan, SOX9 and pSOX9 and that PDGF-B increases the expression of aggrecan in a SOX9-dependent manner. Protein expression was first normalized to actin (internal control), and then compared with expression in pVICs without PDGF-B treatment. Each dots represents an independent experiment. n=4 replicates. (E, F) qRT-PCR for the expressions of candidate ECM degradation enzymes. RNA was collected from pVICs treated with vehicle or 100 ng/ml of PDGF-B for 24 hours in the presence (E) or absence (F) of *Sox9* siRNA. Dotted line represents expression in vehicle treated cells. Each dots represents an independent experiment. N=5-6 replicates. Statistical significance was assessed using a two-tailed, unpaired *t* test in (A and B); one-way ANOVA and Tukey’s multiple comparisons test in (D); and one sample *t* and Wilcoxon test in (E and F). Each dot represents the average from 4-6 staining sections per mouse valve (B and D). Data is represented as Mean ± SEM in (A and B) and Mean ± SD in (E and F). Each dots represent an independent experiment. N=6 in (E) n=5 in (F). Statistical significance was assessed using * = *p*<0.05. ** = *p*<0.01. *** = *p*<0.001.

SOX9 activity is regulated by several post-translational mechanisms, which includes phosphorylation ^43^. To investigate the relationship between PDGF-B signaling, SOX9 and proteoglycan expression, we treated porcine aortic valve VICs (pVICs) with the growth factor PDGF-B, and observed increased SOX9, phospho-SOX9 and aggrecan expression (Figure 6C, D). Additionally, PDGF-B differentially regulated the expression of ADAMTS proteases in pVICs: *Adamts9* and *Adamts5* were downregulated by PDGF-B treatment, whereas *Adamts4* and *Adamts1* were upregulated (Figure 6E). Furthermore, PDGF-B-mediated changes in the levels of aggrecan and *Adamts5* in pVICs were dependent on the expression of SOX9. Specifically, PDGF-B failed to upregulate aggrecan and downregulate *Adamts5* in siSox9-transfected pVICs (Figure 6C, D, F). These results suggest that PDGF-B signaling upregulates proteoglycan expression in a SOX9-dependent mechanism.

### PDGFB and SOX9 are elevated in the myxomatous valves of patients

To investigate the clinical relevance of our findings, we examined whether the phenotype of *Prox1^ΔVEC^* mice recapitulates human valve disease. Myxomatous degeneration is the most common pathology associated with MVP in humans. Recently, the expression of VEC, VIC and ECM molecules was evaluated in the valve tissue collected from MVP patients ^44^. Using these samples (Supplemental Table 1), we found that the expression of the proteoglycans Lumican and *Versican* negatively correlated with *PROX1* (Figure 7A). Moreover, *PDGFB* positively correlated with *SOX9* and *Versican* (Figure 7B).

**Figure 7:**
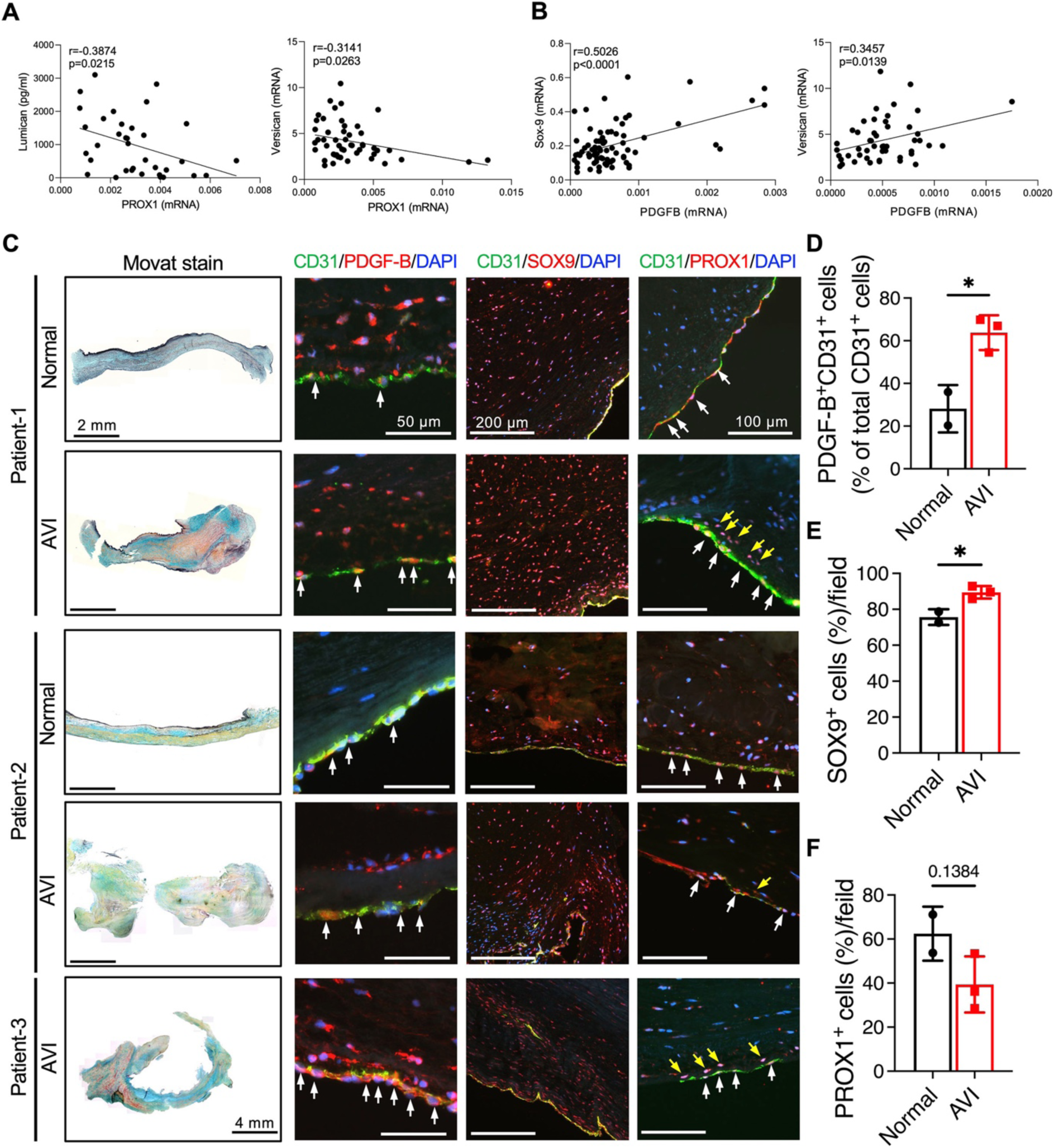
PDGF-B and SOX9 are increased in the myxomatous valves from human patients. (A, B) Protein and RNA samples were obtained from the mitral valves of MVP patients (n=77) who underwent mitral valve replacement surgery. Genes and protein expressions were quantified by qRT-PCR and ELISA respectively. Pearson correlation method was used for calculating r and p. (C) Aortic valve leaflets from 3 patients with aortic valve insufficiency were histologically analyzed. The relatively normal-looking leaflet was used as the internal control. The normal and pathological leaflets were analyzed using Movat Pentachrome stain and by immunohistochemistry for PDGF-B, SOX9, and PROX1. (D-F) Expression of PDGF-B and SOX9 was increased in the diseased leaflet, but no significant difference was observed in the number of PROX1^+^ cells. In PROX1 panels white arrows indicate PROX1^+^ VECs and yellow arrows indicate PROX1^+^ cells of unknown identity in the valve interstitium. Data is represented as Mean ± SEM. * = *p*<0.05. Statistical significance was assessed using a two-tailed, unpaired *t* test. Quantification was performed by averaging the data from 5 sections. PDGF-B staining was quantified as a percentage of PDGFB^+^CD31^+^cells to total CD31^+^ cells. Abbreviations: AVI, leaflet with aortic valve insufficiency.

We analyzed aortic valve samples from three patients (5-year-old female, 7-year-old male and 20-year-old male) with aortic valve stenosis and regurgitation. All patients had congenital aortic stenosis which was initially treated with balloon valvuloplasty. The patients subsequently developed insufficiency which required aortic valve replacement. Valve tissue from two patients had a normal-looking leaflet and a pathological leaflet. Valve tissue from the third patient did not have any normal-looking leaflet. All the leaflets were histologically analyzed.

The H&E and Movat Pentachrome-staining revealed that the pathological valve leaflets showed increased proteoglycan levels (blue) and increased thickness compared with the normal valve leaflet (Figure 7C), consistent with myxomatous degeneration. Additionally, the pathological leaflets showed elevated PDGF-B expression and higher numbers of SOX9^+^ cells relative to the normal leaflet (Figure 7C-E). However, the diseased aortic valve showed normal PROX1 expression in VECs (Figure 7C, white arrows and 7F). Additionally, ectopic CD31^-^ PROX1^+^ cells of unknown identity were observed in the diseased tissue (Figure 7C, yellow arrows). These results suggest that PDGFB-PDGFRβ signaling is upregulated in myxomatous valves irrespective of the etiology.

### Imatinib attenuates aortic valve degeneration in *Prox1^ΔVEC^* mice

Imatinib is an FDA approved drug that inhibits the activity of RTKs such as PDGFRα, PDGFRβ, c-Kit and BCR-ABL ^45^. Although imatinib is not specific for PDGFRβ, it has an excellent safety record in both humans and mice and it is used to treat leukemias and gastrointestinal cancer in humans. Most relevant to this work, imatinib ameliorated connective tissue disorders in patients with PDGFRβ hyperactivating mutations ^35^.

Given our data suggesting that loss of PROX1 in heart valves activates PDGF-B signaling, we considered that imatinib could be repurposed to inhibit the onset and progression of valve defects in *Prox1^ΔVEC^* mice. To test this hypothesis, starting at postnatal day (P) 30 we orally administered imatinib to control and *Prox1^ΔVEC^* mice daily at a dose of 50 mg/kg for 6 months (Figure 8A). Echocardiography showed that imatinib treatment preserves aortic valve function in *Prox1^ΔVEC^* mice by maintaining cusp separation and preventing an increase in peak velocity (Figure 8B). We also observed platelet aggregate-like structures in untreated mutant aortic valves, which were reduced by imatinib treatment (Figure 8C). Histological data suggested that imatinib treatment does not ameliorate the progressive thickening of *Prox1^ΔVEC^* valves (Figure 8C). However, imatinib significantly reduced the abnormal elastin expression on the downstream side of aortic valves in *Prox1^ΔVEC^* mice (Figure 8D, orange arrowhead) while maintaining its expression on the upstream side (Figure 8D, green arrowhead). Imatinib also reduced the expression of versican in the aortic valves of *Prox1^ΔVEC^* mice (Figure 8E). Similarly, although imatinib did not reduce the thickness of the mitral valves it significantly normalized elastin and versican expression in *Prox1^ΔVEC^* mice (Figure S18). Furthermore, imatinib normalized the expression of SOX9 in the aortic and mitral valves of *Prox1^ΔVEC^* mice (Figure S19). These results reveal that imatinib treatment can partially rescue the valve defects of *Prox1^ΔVEC^* mice by normalizing the expression of SOX9 and ECM components.

**Figure 8:**
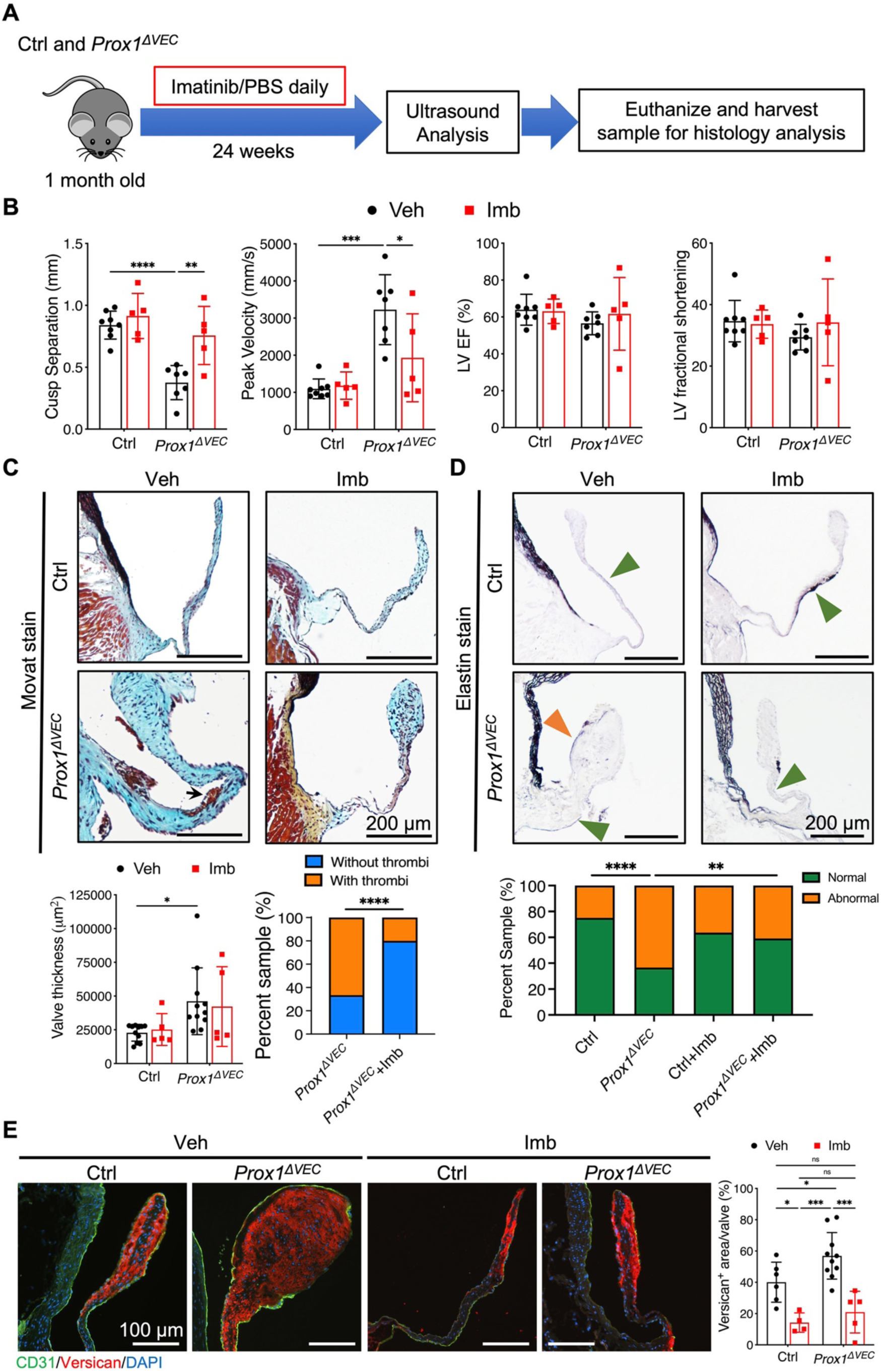
RTK inhibitor imatinib partially rescues aortic valve function in *Prox1^ΔVEC^* mice. (A) Schematic of the experimental design. Imatinib (50mg/kg-BW/day) or PBS (vehicle) was orally administrated to control and *Prox1^ΔVEC^* mice daily for 24 weeks starting from when they were 1-month-old. At the end of the treatment aortic valves were evaluated by echocardiography and histology. (B) Echocardiography showed that imatinib enhanced cusp separation and lowered aortic peak velocity in *Prox1^ΔVEC^* mice when compared to untreated *Prox1^ΔVEC^*mice. There were no significant differences in left ventricle functions (ejection fraction and fractional shortening). N=7 vehicle treated controls, n= 8 vehicle treated *Prox1^ΔVEC^* mice, n=5 imatinib treated controls and n=5 imatinib treated *Prox1^ΔVEC^* mice. (C) Representative Movat Pentachrome-stained sections from the aortic valves of control and *Prox1^ΔVEC^* mice treated with vehicle or imatinib. The graphs show the quantification of valve thickness and the presence of thrombi. Arrow points to a thrombus-like structure in an untreated *Prox1^ΔVEC^* mouse. The number of thrombus-like structures was reduced in *Prox1^ΔVEC^* mice treated with imatinib. The area of valve leaflets was measured from 4-6 sections per valve. The average of these areas is represented as the thickness of valve in that mouse. Each dot represents an individual mouse. Valve thickness in *Prox1^ΔVEC^* mice was not reduced by imatinib treatment. N=12 vehicle treated controls, n=10 vehicle treated *Prox1^ΔVEC^* mice, n=5 imatinib treated controls and n=5 imatinib treated *Prox1^ΔVEC^*mice. (D) Abnormal elastin distribution was rescued in the aortic valves of imatinib treated *Prox1^ΔVEC^* mice. N=10 vehicle treated control, n=10 vehicle treated *Prox1^ΔVEC^*, n=5 imatinib treated control and n=5 imatinib treated *Prox1^ΔVEC^* mice. (E) IHC and quantification indicated that abnormal versican production in *Prox1^ΔVEC^* mice was inhibited by imatinib treated. N=6 vehicle treated controls, n=10 vehicle treated *Prox1^ΔVEC^* mice, n=4 imatinib treated controls and n=5 imatinib treated *Prox1^ΔVEC^* mice. Data is represented in Mean ± SD in (B) and Mean ± SEM in (C and E). Statistical significance was assessed using a two-way ANOVA and Tukey’s multiple comparisons test in (B, C, and E) and using the Fisher exact test in (C and D). * = *p*<0.05; ** = *p*<0.01, *** = *p*<0.001, **** = *p*<0.0001 and. Each dot represents the average of 3-5 sections from a mouse. Abbreviations: Veh, vehicle; Imb, imatinib.

In summary, we have shown that PROX1 and FOXC2, molecules that are critical for the development of lymphatic vasculature and vascular valves, are necessary to prevent the myxomatous degeneration of heart valves.

## DISCUSSION

Myxomatous degeneration causes MVP, one of the more common valve disorders, affecting about 144 million individuals worldwide ^46^. MVP predisposes to mitral regurgitation, pulmonary hypertension, left ventricular hypertrophy, arrhythmias, endocarditis, heart failure and sudden death. MVP takes decades to progress from the time of diagnosis to the time when it requires surgical intervention. Therefore, prophylactic drugs can potentially prevent the progression of MVP.

Here, we describe a new mouse model for myxomatous degeneration of heart valves and reveal that PROX1 and its target FOXC2 are required to prevent valve thickening and dysfunction in both male and female mice. Our data support a model in which PDGF-B expression is elevated in the absence of PROX1 or FOXC2. PDGF-B from VECs activates PDGFRβ in VICs to upregulate SOX9, downregulate ADAMTS and ultimately leading to increased proteoglycan levels (Figure S20). These findings were reinforced in a human cohort of MVP valves, showing an inverse correlation between PROX1 and proteoglycan expression. Conversely, PDGF-B expression positively correlated with SOX9 and Versican.

PDGF-B is required for the development of heart valves, as *Pdgfb^−/−^*embryos have hypoplastic heart valves ^31^. Mutations that constitutively activate PDGFRβ have been identified in Kosaki overgrowth syndrome patients with MVP and pulmonary valve stenosis ^33,35^. Our data show that aberrant activation of PDGFRβ signaling is associated with myxomatous degeneration in mice. Furthermore, the RTK inhibitor imatinib largely ameliorates the heart valve defects of *Prox1^ΔVEC^* mice. Thus, our study suggests that the hyperactivation of PDGFB/ PDGFRβ could lead to MVP and the progression of this disease can potentially be slowed down by imatinib, a drug with a strong record of patient safety.

PDGF-B is required to recruit mural cells to the collecting lymphatic vessels that express PROX1 at low levels ^47^. In contrast, PROX1^high^ lymphatic capillaries and lymphatic valves lack PDGF-B expression and mural cell coverage ^37^. Lymphatic capillaries of mice lacking PROX1 abnormally recruit PDGFRβ^+^ mural cells ^48^, suggesting that PROX1 might be inhibiting PDGF-B expression in the lymphatic vasculature. Thus, PDGF-B is a potential target of PROX1 in both lymphatic endothelial cells and VECs. PDGF-B is also expressed in the PROX1^-^ VICs of *Prox1^ΔVEC^* mice. PDGF-B could promote its own expression in certain cell types ^49,50^. Thus, we speculate that PDGF-B from VECs promotes *Pdgfb* expression in the VIC of *Prox1^ΔVEC^* mice.

The mechanisms by which PROX1 and FOXC2 regulate PDGF-B expression in VECs are currently unknown. PDGF-B is regulated by shear stress in vascular endothelial cells ^51^. PROX1 and FOXC2 regulate cytoskeletal organization and shear stress response in lymphatic endothelial cells ^11,52^. Therefore, it is possible that PROX1 and/or FOXC2 directly or indirectly regulate PDGF-B expression in VECs in response to shear stress.

PDGF-B is considered as the canonical ligand of PDGFRβ. However, PDGFRα is considered as a receptor for both PDGF-A and PDGF-C. Therefore, the potential interaction between PDGF-B and PDGFRα cannot be excluded in certain tissues at specific time points. Important for this work, PDGFRα was recently reported to be expressed in downstream VECs ^53^. Conditional deletion of *Pdgfra* from VECs resulted in the disruption of the endothelial cell layer and early onset myxomatous degeneration with increased pERK1/2 in the anterior leaflet of mitral valves. Although our data suggests that PROX1 functions by antagonizing the PDGF-B/PDGFRβ pathway, it will be important to explore the relationship between PROX1, PDGF-B and PDGFRα.

Limitations of this study: FOXC2 and PDGF-B may not be the only targets of PROX1 in VECs. FOXC2 does not appear to play a significant role in mitral VECs. Furthermore, mutations in genes that regulate TGFβ signaling, such as *TGFBR1, TGFBR2, SMAD2, SMAD3* and *FBN1*, are associated with MVP ^54^. The inability of imatinib to prevent the thickening of *Prox1^ΔVEC^* valves could be due to the upregulation of *TGFB1* in these tissues (Figure 4A). Further research is required to identify the relationship between PROX1, *TGFB1* and other modulators of valve disease.

## METHODS

### Mice

*Notch-Cre^Lo^*, *Tie2-Cre*, *Rosa26^mT/mG^*, *Rosa26^+/tdTom^* and CAG-LoxP-Stop-LoxP-rtTA3-mKate2 were purchased from Jackson Laboratory (stock no. 006953; 008863; 007676; 007909 and 029633 respectively) *Prox1-Tom* (stock no. 036531-UCD) was purchased from MMRRC. *Prox1^+/-^*, *Prox1^+/flox^*, *Pdgfrb^+/LSL-D849V^*, NFATC1^enCre^ and *Rosa26^+/LSL-PDGFB^* mice were reported previously ^13,14,32,36,55^. *Prox1-2A-Cre* and shFoxc2 models were generated by Cyagen Inc and Mirimus Inc respectively on a fee-for-service basis. *Prox1^+/-^* mice were maintained in NMRI background. The rest of the mice were maintained in mixed background. Both males and virgin females were used in all analysis. All mouse experiments are followed US National Institutes of Health guidelines (Guide for the Care and Use of Laboratory Animals) and were performed with the protocols approved by the OMRF’s IACUC approval #21-20.

### Embryo and Tissue Collection

Adult mice were euthanized by asphyxiation with CO_2_ followed by cervical dislocation. Embryos were quickly removed from the uterus and anesthetized using ice-cold phosphate-buffered saline (PBS). Thoracotomy was performed to facilitate the drainage of blood from the embryos. Subsequently, embryos were analyzed by whole-mount IHC or cryo-IHC as described below.

Adult mice were asphyxiated and perfused with ice-cold PBS and followed by 4% ice-cold PFA. Subsequently, hearts were processed for cryosections, paraffin sections, SEM or TEM as described below in a double-blinded manner.

### IHC of paraffin sections

Paraffin-embedded tissue sections were de-paraffinized, rehydrated and subjected to antigen retrieval by boiling in pH 6.0 or pH 9.0 Tris-EDTA buffer (1mM EDTA, 10mM Tris, 0.05% Tween20) for 30 minutes in a steamer. Antigen-retrieved sections were blocked using 5% donkey serum (Sigma; D9663) or 5% bovine serum albumin (BSA: AmericanBIO; ABO1088) in PBST (PBS + 0.1% Triton X-100) for 1 hour at RT, incubated with primary antibodies diluted in the blocking solution overnight at 4°C. On the next day the sections were incubated with fluorescent secondary antibodies diluted in the blocking solution for 2 hours at RT.

pH 6.0 Tris-EDTA buffer was used as the antigen-retrieval buffer for the following antibodies: CD31 (R&D systems; AF3628; 1:500), Aggrecan (Millipore; AB1031; 1:200), PROX1 (R&D systems; AF2727; 1:200), and DPEAAE (cleaved-versican, V0, V1 Neo) (Invitrogen; PAI-1748A; 1:400).

pH 9.0 Tris-EDTA buffer was used as the antigen-retrieval buffer for the following antibodies: CD31 (R&D systems; AF3628; 1:500), SOX9 (Millipore; AB5535; 1:500), PDGF-BB (Abcam; ab23914; 1:200), and Collagen Hybridizing Peptide, cy3 conjugate. (3Helix; R-CHP; 5 μM).

To detect Versican the sections were antigen-retrieved using pH 9.0 Tris-EDTA buffer, treated with chondroitinase ABC (Sigma; C2905) for 1 hour at 37°C before incubating with the primary antibody (Millipore; AB1033; 1:300) overnight at 4°C.

Negative controls (no primary antibody) were used for all experiments. On the next day, sections were washed 3X in PBST and incubated with fluorescence-conjugated secondary antibodies for 2 hours at RT, washed 3X with PBST and mounted with Fluoro Gel with DABCO anti fading mounting medium (Electron Microscopy Sciences; 17985-03) with DAPI (4’, 6-Diamidino-2-Phenylindole) (Life Technologies; D1306; 1:10,000). Images were captured using a Nikon Eclipse 80i system with the NIS-Elements BR software (version 4.3) or Nikon C2 confocal microscope equipped with a motorized stage with the NIS-element AR software (version 5.02).

### IHC of cryosections

Embryos and hearts were fixed overnight in 2% PFA at 4°C. After washing profusely in ice-cold PBS the tissues were immersed in 15% sucrose overnight at 4°C and then in 30% sucrose overnight at 4°C, cryoembedded in OCT compound (Sakura; 4583) and stored at −80°C. Cryosections (10 µm) were collected using a cryotome (Thermo HM525NX) and directly mounted onto positively charged slides. Slides were air-dried for 30 minutes at 60°C, washed with PBS to remove OCT, blocked using 5% donkey serum or 5% BSA in PBST for 1 hour at RT and incubated overnight at 4°C with primary antibodies diluted in the blocking solution: CD31 (R&D systems; AF3628; 1:500), Aggrecan (Millipore; AB1031; 1:200), PROX1 (R&D systems; AF2727; 1:200), PROX1 (AngioBio; 11-002; 1:200), VEGFR3, (R&D systems; AF743; 1:400) and GFP (Abcam; ab13970; 1:1000). On the next day, slides were incubated with fluorescence-conjugated secondary antibodies, mounted and imaged as described above.

### Whole-mount IHC

Mesenteries were harvested from freshly collected embryos, fixed in 2% paraformaldehyde overnight at 4°C, washed profusely in ice-cold PBS, and then used for whole-mount IHC. Embryos were fixed in 2% paraformaldehyde overnight at 4°C, washed profusely in ice-cold PBS and then the dermal skin samples were isolated for whole-mount IHC. A detailed description of the whole-mount IHC protocol was recently published ^56^. Briefly, the samples were washed with PBST (0.2% Triton X-100 in PBS), sequentially incubated overnight at room temperature in PBSTD (0.2% Triton X-100, 20% DMSO in PBS), PBSTTDND (0.1% Tween X-100, 0.1% Triton X-100, 0.1% deoxycholate, 0.1% NP40, 20% DMSO in PBS) and PBSTDM (0.2% Triton X-100, 20% DMSO, 0.3 M glycine in PBS). The pretreated samples were blocked with PBSTD (0.2% Triton X-100/10% DMSO/6% donkey serum in PBS) overnight, washed 2X in PTwH (0.2% Tween-20, 10 µg/mL heparin, 10% DMSO), and incubated overnight with primary antibodies diluted in PTwHD (0.2% Tween-20, 10 µg/mL heparin, 10% DMSO, 3% donkey serum in PBS) at RT. After profuse washing in PTwH the samples were incubated overnight with secondary antibodies diluted in PTwHD (0.2% Tween-20, 10 µg/mL heparin, 3% donkey serum in PBS). Samples were washed again in PTwH, mounted using homemade mounting media and visualized using Nikon Eclipse 80i system with NIS-Elements AR software (version 3.0).

### Image Analysis

Image analysis were performed by using ImageJ2 (Fiji; Version 2.3.0/1.53qr). Quantification was performed by measuring total number of positive-pixels or positive-cells in the whole mouse valve leaflet. For the human valve sample 5 images at 10x magnification were acquired per valve.

### ECM analysis

Hearts were isolated and fixed in 4% paraformaldehyde (PFA) overnight at 4°C, dehydrated with graded ethanol series, cleared in xylene, embedded in paraffin and sectioned at 5 μm thickness. Movat pentachrome (Abcam; ab245884), Alizarin red (IHC World; IW-3001) and Resorcin-Fuchsin (Electron Microscopy Sciences; 26370-01) staining’s were performed using the manufacturer’s protocols. Brightfield images were taken by using Nikon Eclipse 80i system with NIS-Elements AR software (version 3.0).

### Valve thickness measurement

4-7 sections at 100-150 μm intervals were collected for each valve. The total area of the valve leaflets (2 leaflets from base to tip) was calculated for each slide and the average area from all the slides was used as the measure of valve thickness.

### Ultrasound Imaging

Valve function was monitored by the recently established, modified-view, echocardiography ultrasound imaging technique using Vevo 2100. This B-mode imaging technique captures high-resolution images of the valves ^57^. Briefly, 1%-3% isoflurane was continuously administered through a nose cone to anesthetize the mice. To ensure accuracy of all measurements, physiological parameters were monitored, and recordings were taken only if the heart rate was 450 to 550 beats per minute and respiration was 50 to 100 breaths per minute. Echocardiography was performed by using B-mode imaging with a Vevo 2100 MS550D transducer (VisualSonics). Distance between aortic valve leaflets, left ventricular outflow tract (LVOT), peak blood flow velocity across the valve, left ventricle ejection fraction, and left ventricle fraction shortening were measured. LVOT was measured and used to normalize the cusp separation and defined as fractional valve opening. We also measured peak blood flow velocity across the valves (jet) with a combination of color and pulse-wave Doppler that allows alignment along the maximum flow direction. Left ventricle function was determined by left ventricle ejection fraction, and left ventricle fraction shortening.

### RNAscope

10 µm thick cryosections of heart samples were prepared as described above. These cryosections were used with RNAscope 2.5 HD Detection Kit-RED (ACDbio; 322360) according to the manufacturer’s instructions.

Cryosections were incubated with RNAscope hydrogen peroxide-reagent (ACDbio; 322330) for 10 minutes at room temperature and then with target retrieval reagent (ACDbio; 322000) for 5 minutes at 95°C. Slides then underwent Protease Plus treatment (ACDbio; 322331) for 30 minutes at 40°C, followed by two distilled water washes for 2 minutes each. Sections were hybridized with RNAscope probes against *Pdgfb* (ACDbio; 424651), positive control probe (ACDbio; 313911), and negative control probe (ACDbio; 310043) for 2 hours at 40 °C, followed by two washes in 1 × RNAscope Washing Buffer (ACDbio; 310091), 2 minutes each. After hybridization, sections were sequentially incubated with 6 amplification buffers: AMP1 (30 minutes, 40°C), AMP2 (15 minutes, 40°C), AMP3 (30 minutes, 40°C), AMP4 (15 minutes, 40°C), AMP5 (30 minutes, RT) and AMP6 (15 minutes, RT). In between the amplification buffers the slides were washed 2X at room temperature with 1 × RNAscope Washing Buffer (2 minutes each). Sections were incubated in Fast RED mix for 4 minutes at room temperature and washed 3X with PBS for 2 minutes each. Lastly, IHC was performed for CD31 and DAPI.

The images were taken by using a Nikon C2 confocal microscope with the NIS-element AR software. The data was calculated as a Histo score (H score) according to the manufacturer’s instructions. First, each cell within an image is assigned a Bin value based on the size and the number of dots (puncta) per cell. Bin0: 0 pucta/cell; Bin1: 1-3 dots/cell; Bin2: 4-9 dots/cell; Bin3: 10-15 dots/cell; Bin4: >15dots/cell). Then, the percentage of cells with Bin values 0-4 was calculated. The H-score was then calculated using the formula

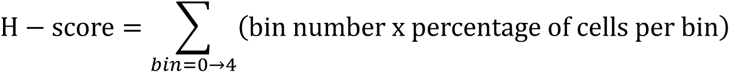

### Scanning Electron Microscopy

SEM was performed as described in detail previously ^7,58^. Heart valve tissue were fixed in a fix solution of 2% PFA and 2.5% glutaraldehyde in 0.1 M sodium cacodylate buffer at 4°C for 24 hours. After washing profusely in PBS, tissues were post fixed in 1% osmium tetroxide in 0.1 M cacodylate buffer for 2 hours at room temperature. Fixed samples were subsequently dehydrated in a graded ethanol series. The tissues were further dehydrated in hexamethldisilazane (HMDS) and air-dried overnight in chemical safety hood. Dried tissues were sputter-coated with Au particles (Med-010 Sputter Coater by Balzers-Union) and observed under JCM-6000PLUS SEM (JEOL-USA) at an accelerating voltage of 15KV.

### Transmission Electron Microscopy

Heart valve tissue were fixed in 2% PFA and 2.5% glutaraldehyde in 0.1 M sodium cacodylate buffer at 4°C for 2 days. After washing several times in PBS, heart valve tissues were embedded in 7% low melting point agarose (Fisher Scientific; BP1360-100), and 1 mm sections were prepared using vibratome (Leica; VT1000S). 1 mm sections that contain the aortic and mitral valves were collected and trimmed into 1 mm cubes to remove other tissues. The cubes were post-fixed in 1% osmium tetroxide for 2 hours and 1% tannic acid for 1 hour. The samples were subsequently dehydrated by gradient ethanol and embedded in epoxy resin (Electron Microscopy Sciences). Semi-thin (500 nm) and ultra-thin (70 nm) sections were obtained using an ultramicrotome (Leica Microsystems; Leica EM UC7) equipped with a diamond knife. Semi-thin sections were stained with Epoxy tissue stain (Electron Microscopy Sciences) and analyzed with an Eclipse E800M microscope (Nikon). Ultrathin sections were stained with uranyl acetate and lead citrate before being viewed with a Hitachi H-7600 electron microscope equipped with a 4-megapixel digital monochrome camera and AMT-EM image acquisition software (Advanced Microscopy Techniques).

### Ovine Mitral Valve Endothelial Cells

Ovine mitral VECs were provided by Dr. Bischoff. Briefly, VEC from mitral valve leaflets of sheep were isolated and prepared as described ^59^. MVECs were expanded and grown on 1% gelatin-coated dishes in endothelial basal medium-2 (EBM-2 (Lonza; 3156) with 10 % heat-inactivated FBS (Gibco), 1X glutamine/penicillin/streptomycin (Life Technologies, Inc) and 2 ng/ml basic fibroblast growth factor (Roche Applied Science). Experiments were performed at passages 6-8.

### Porcine Aortic Valve Interstitial Cells

Porcine VICs were isolated from aortic valve leaflets of pigs as described ^60^. pVICs were isolated, expanded in DMEM (Corning; 10-013-CV) with 10 % heat-inactivated FBS (Gibco), and 1X glutamine/penicillin/streptomycin (Life Technologies, Inc). Experiments were performed at passages 3-5.

### Cell Treatments

#### siRNA transfection

Sheep MVECs and AVICs were seeded onto 6-well plates or 6-cm culture dishes. After 24-36 hours of culture (around 50% confluence), cells were transfected with siControl (Integrated DNA Technologies; 51-01-14-04), siPROX1 for sheep (Invitrogen; 10620310), or siSOX9 for porcine (Life Technologies, Inc; Silencer Select 555768) using Lipofectamine RNAiMax Transfection Reagent (Invitrogen; 13778150) and Opti-MEM + GlutaMax (Gibco; 51986-034) according to manufacturer’s instruction.

#### PDGF-B treatment

Confluent porcine VICs were treated with the indicated concentrations of recombinant human PDGF-BB (PeproTech; 100-14B) and collected after 24 hours for western blotting or qRT-PCR.

### RNA isolation and quantitative PCR

Total RNA was isolated from sheep MVECs, porcine AVICs or mouse heart valve tissue by using Trizol (Invitrogen) and purified by using RNeasy Mini Kit (QIAGEN; 74106) according to manufacturer’s instructions. For mouse tissue, the tissue was cut into small pieces and homogenized by QIAshredder (QIAGEN; 79656) prior to the addition of Trizol. cDNA was synthesized from total RNA (1-2 μg) with iScript Advanced cDNA Synthesis Kit (Bio-Rad; 1725038). Quantitative RT-PCR (qPCR) was performed using SsoAdvanced Universal SYBR Green SuperMix reagent (Bio-Rad; 1725274) in a CFX96 Real-Time System (Bio-Rad). Expression levels were normalized to GAPDH. Primer sequences are provided below.

### Western blot

After siRNA and/or PDGF-BB treatment, porcine VICs were washed quickly with cold PBS and collected by lysis buffer (25 mM Tris pH 7.4, 150 mM NaCl, 0.5% Na deoxycholate, 2% NP-40, 0.2% SDS, 7% glycerol) containing protease and phosphatase inhibitors (Thermo Scientific; 1861281). Cells were scraped into a 1.5-mL tube, lysed on ice for 30 minutes and centrifuged to collect the supernatant as total protein extract. Protein concentrations were determined by BCA protein assay (Thermo Scientific; 23227). 20 μg of protein were loaded to SDS polyacrylamide gel electrophoresis and proteins were transferred to PVDF membrane using standard wet transfer method (80 volt, 3 hours on ice). The blots were probed with indicated primary antibodies listed below. For loading control, blots were subsequently probed with pan-actin. The protein bands were quantified by Image J and normalized to the corresponding loading controls.

### Imatinib Treatment

Control and *Prox1^ΔVEC^* mice were randomized into 2 groups (vehicle and Imatinib). Mice were gavaged every day with Imatinib (50 mg/kg-BW/day; Cayman; 13139) or PBS by using plastic feeding tube (20 ga x 38 mm; INSTECH; FTP-20-38) for 24 weeks starting from when they were 1-month-old. After 24 weeks of treatment valve function was analyzed by echocardiography. Subsequently, the mice were sacrificed, and their valves were harvested and analyzed by histology.

### Human valve tissues

The human MVP samples were recently reported (Table S1) ^44^. Using the cDNA samples reported in the original publication the expression of *PROX1, PDGFB, Versican* and *SOX9* was analyzed by qRT-PCR. The expression of Lumican was analyzed by ELISA using the protein lysates that were collected for the original publication.

The three insufficient aortic valves were donated by deidentified patients undergoing aortic valve replacement surgery at the Oklahoma Children’s Hospital. The patients were a 5-year-old female, 7-year-old male and a 20-year-old male. All patients were evaluated by transthoracic echocardiography. Informed consent was obtained from each patient and control and the study protocol conforms to the ethical guidelines of the 1975 Declaration of Helsinki as reflected in a priori approval by the institution’s human research committee.

### Statistics

Statistical analyses were performed by using GraphPad Prism 9 (version 9.4.1). Details for specific statistical analyses can be found in the figure legends. For parametric comparison, the two-tailed, unpaired *t* test was used to compare means between two experimental groups. For comparison of percentage from two groups, the Fisher exact test was performed. For non-parametric comparisons, Mann-Whitney *U* tests were used. Multiple means were compared using a one-way ANOVA and Tukey’s multiple comparisons test, and two-way ANOVA and Tukey’s multiple comparisons test was used to compare several treatment groups examined with 2 or more independent variables. For small samples comparison, we used one sample *t* and Wilcoxon test. A *P* value <0.05 was considered statistically significant and is indicated as \**P*<0.05, \*\**P*<0.01, \*\*\**P*<0.001, **** *P*<0.0001.

### List of antibodies

| Antibody | Vender | Cat No: | Dilution | Note |
| --- | --- | --- | --- | --- |
| Aggrecan | Millipore | AB1031 | 1:200 | Paraffin section<br>Cryo section<br>Western blotting<br>(non-reducing protein) |
| CD31 | R&D systems | AF3628 | 1:500 | Paraffin section<br>Cryo section |
| Collagen Hybridizing<br>Peptide Cy3-conjugated | 3Helix | R-CHP | 5μM | Paraffin section<br>Cryo section |
| DPEAAE (cleaved-versican,<br>V0, V1 Neo) | Invitrogen | PAI-1748A | 1:400 | Paraffin section<br>Cryo section |
| FOXC2 | Santa Cruz | Sc-515234 | 1:500 | Paraffin section<br>Cryo section |
| GFP | Abcam | ab13970 | 1:1000 | Paraffin section<br>Cryo section<br>Whole-mount staining |
| Notch1 | Cell Signaling | 3608 | 1:1000 | Western blotting |
| Cleaved Notch1 | Cell Signaling | 4147 | 1:500 | Western blotting |
| PDGF-BB | Abcam | ab23914 | 1:200 | Paraffin section<br>Cryo section |
| PROX1 | R&D systems | AF2727 | 1:200 | Paraffin section<br>Cryo section<br>Western blotting |
| PROX1 | AngioBio | 11-002 | 1:200 | Cryo section<br>Whole-mount staining |
| SOX9 | Millipore | AB5535 | 1:500 | Paraffin section<br>Western blotting |
| von Willebrand Factor | Dako | A0082 | 1:500 | Paraffin section<br>Cryo section |
| Anti-Actin, α-Smooth<br>muscle-Cy3 | Sigma | C6198 | 1:3000 | Paraffin section<br>Cryo section |
| Phospho-SOX9 (S181) | Abcam | Ab59252 | 1:500 | Western blotting |
| VEGFR3 | R&D systems | AF743 | 1:400 | Cryo section<br>Whole-mount staining |
| Versican | Millipore | AB1033 | 1:300 | Paraffin section<br>Cryo section |
| β-Actin | Sigma | A5441 | 1:100000 | Western blotting |
| Donkey anti-Goat Cy3-conjugated | Jackson Immuno Research | 705-165-147 | 1:500 | Secondary antibody for IF |
| Donkey anti-Rabbit Cy3-conjugated | Jackson Immuno Research | 711-165-152 | 1:500 | Secondary antibody for IF |
| Donkey anti-Goat Cy5-conjugated | Jackson Immuno Research | 705-175-147 | 1:500 | Secondary antibody for IF |
| Donkey anti-Chicken FITC-conjugated | Jackson Immuno Research | 703-095-155 | 1:500 | Secondary antibody for IF |
| Donkey anti-Rabbit Alexa Fluor 488-conjugated | Jackson Immuno Research | 711-547-003 | 1:500 | Secondary antibody for IF |
| Donkey anti-Goat Alexa Fluor 488-conjugated | Jackson Immuno Research | 705-545-147 | 1:500 | Secondary antibody for IF |
| Donkey anti-Chicken Alexa Fluor 488-conjugated | Jackson Immuno Research | 703-545-155 | 1:500 | Secondary antibody for IF |
| Donkey anti-Rabbit Alexa Fluor 647-conjugated | Jackson Immuno Research | 711-605-152 | 1:500 | Secondary antibody for IF |
| Anti-Mouse HRP-conjugated | Sigma | A4416 | 1:5000 | Secondary antibody for WB |
| Goat anti-Rabbit, HRP-conjugated | Immuno-Reagents | GtxRb-003-EHRPX | 1:5000 | Secondary antibody for WB |
| Donkey anti-Goat HRP-conjugated | Jackson Immuno Research | 705-035-003 | 1:5000 | Secondary antibody for WB |

### List of Primers for qPCR

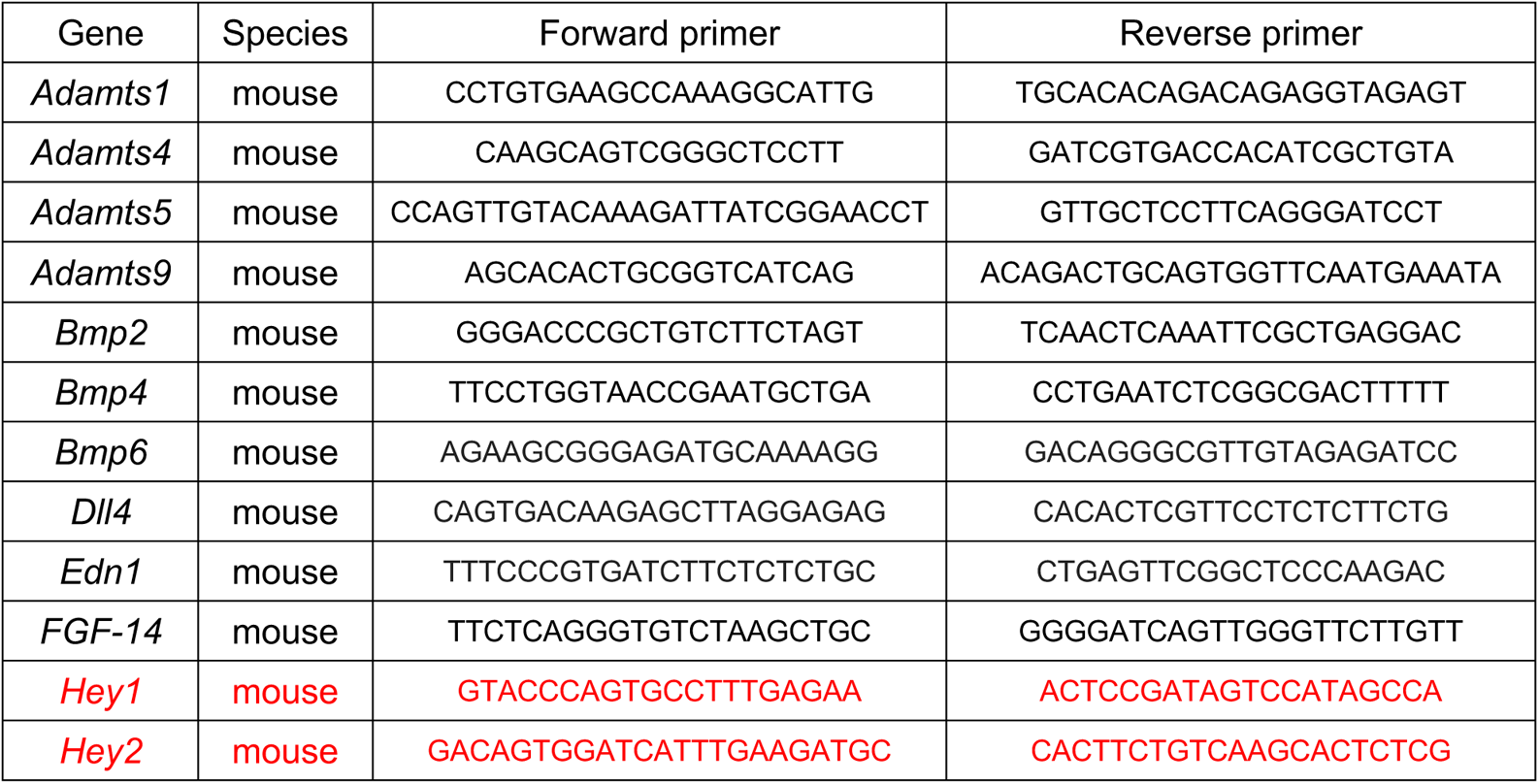
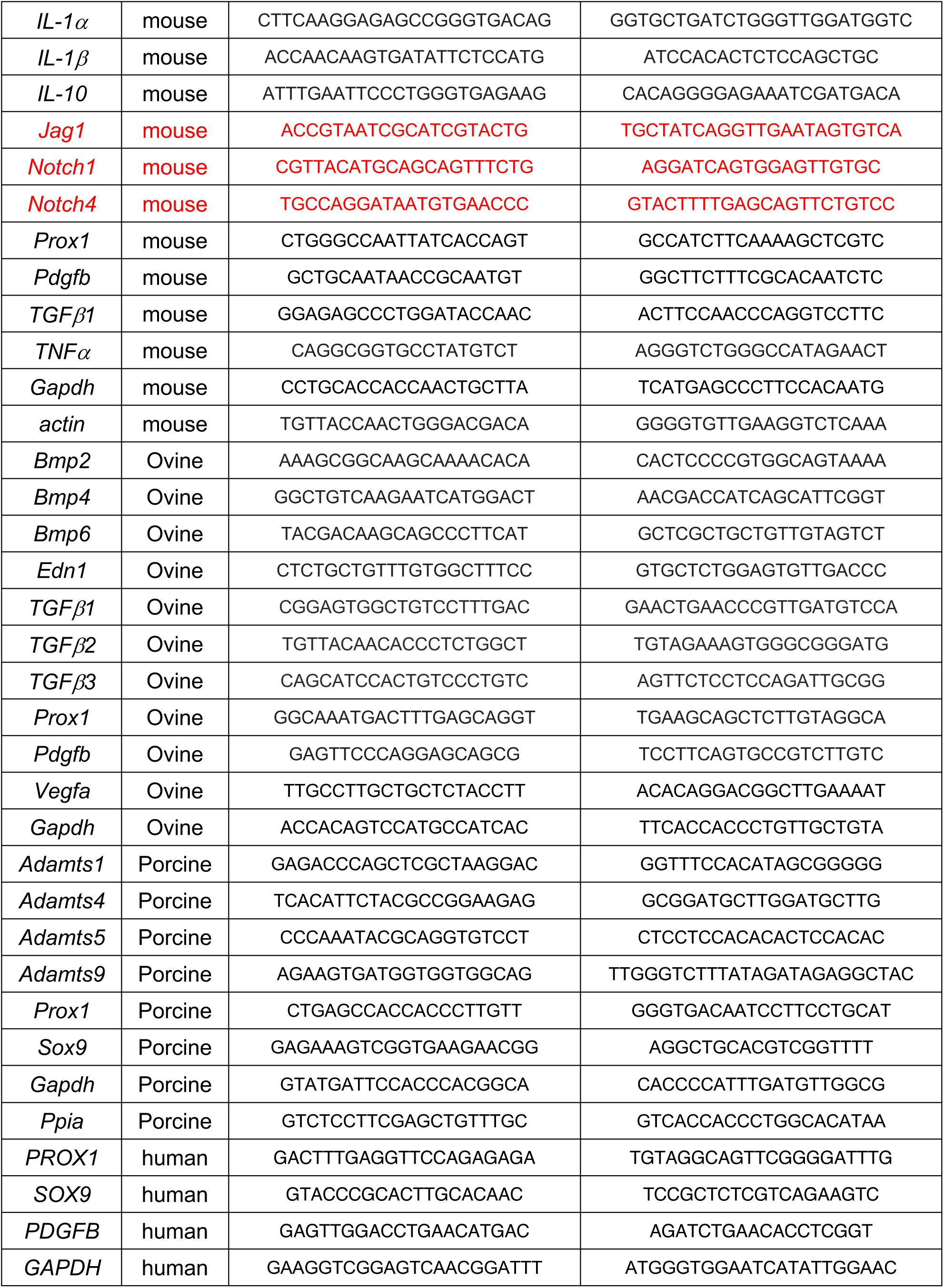

## Affiliations

Cardiovascular Biology Research Program, Oklahoma Medical Research Foundation, 825 NE 13^th^ street, Oklahoma City, OK 73104, USA (YCH, XG, RV, KS, ZA-N, JK, RS, LC, BC, LX, FL, LO, JA, RSS); Division of Molecular Cardiovascular Biology, Cincinnati Children’s Hospital Medical Center, 3333 Burnet Ave, Cincinnati, OH 45229, USA (AO, KEY); Molecular Cardiology Research Institute, Tufts Medical Center, Boston, MA (JI); Cardiovascular Translational Research, Navarrabiomed (Miguel Servet Foundation), Instituto de Investigación Sanitaria de Navarra (IdiSNA), Hospital Universitario de Navarra (HUN), Universidad Pública de Navarra (UPNA), Pamplona, Spain (JI, AF-C, NL-A); Department of Cell Biology, University of Oklahoma Health Sciences Center, Oklahoma City, OK 73117, USA (JK, LO, RSS); Vascular Biology Program, Boston Children’s Hospital, Boston, MA 02115, USA (HC, JB); Department of Genetics, Albert Einstein College of Medicine, Bronx, NY 10461, USA (BZ); Oklahoma Children’s Hospital, University of Oklahoma Health Heart Center, Oklahoma City, OK 73104, USA (HMB); Department of Medicine, Cardiovascular Division, Brigham and Women’s Hospital, Boston, MA 02115, USA (EA); Sanegene Bio, 400 TradeCenter, Woburn, MA, 01801, USA (XG); Daegu Gyeongbuk Medical Innovation Foundation, Daegu 41061, Republic of Korea (BC).

## Acknowledgements

We thank Dr. Christer Betsholtz for sharing the *R26^+/LSL-PDGFB^* mouse line, Dr. Fumio Matsuzaki for the *Prox1^+/flox^*line, Dr. Wei Jing for help with data organization and Dr. Angela Andersen, Life Science Editors, for editing the manuscript.

## Author Contributions

YCH, XG, RV, KS, ZA-N, RS-M, LC, BC, JI, AF-C, NL-A and RSS performed experiments and analyzed data. JK, JW-S, ZA, HB, LEO and JB provided critical reagents; HB, HC, LX, FL, EA, LEO, JA and JB provided input; YCH and RSS co-wrote the manuscript with input from all authors. RSS conceived the project and obtained funding.

## Data Availability

No datasets were generated for this work.

